# Forest growth resistance and resilience to the 2018-2020 drought depend on tree diversity and mycorrhizal type

**DOI:** 10.1101/2024.01.23.576797

**Authors:** Lena Sachsenmaier, Florian Schnabel, Peter Dietrich, Nico Eisenhauer, Olga Ferlian, Julius Quosh, Ronny Richter, Christian Wirth

## Abstract

1. The frequency of consecutive drought years is predicted to increase due to climate change. These droughts have strong negative impacts on forest ecosystems. Mixing tree species is proposed to increase the drought resistance and resilience of tree communities. However, this promising diversity effect has not yet been investigated under extreme drought conditions and in the context of complementary mycorrhizal associations and their potential role in enhancing water uptake.
2. Here, we investigate whether tree diversity promotes growth resistance and resilience to extreme drought and whether drought responses are modulated by mycorrhizal associations. We used inventory data (2015 – 2021) of a young tree diversity experiment in Germany, manipulating tree species richness (1, 2, 4 species) and mycorrhizal type (communities containing arbuscular mycorrhizal (AM) or ectomycorrhizal (EM) tree species, or both). For all tree communities, we calculated basal area increment (BAI) in the periods before, during, and after the drought and used the concepts of resistance and resilience to quantify growth responses to drought.
3. We found strong growth declines during the extreme 2018-2020 drought for most of the tree communities. Contrary to our hypothesis, we did not find that tree species richness *per se* can buffer the negative impacts of extreme drought on tree growth. However, while for EM communities, drought resistance, and resilience decreased with tree species richness, they increased for AM communities and communities comprising both mycorrhizal types. We highlight that among various tree species mixtures, only those with mixed mycorrhizal types outperformed their respective monocultures during and after drought. Further, under extreme drought, the community tends to segregate into “winner” and “loser” tree species in terms of diversity, indicating a potential intensification of competition.
4. While we cannot disentangle the underlying mechanisms, or clarify the role of mycorrhiza during drought, our findings suggest that mixtures of mycorrhizal types within tree communities could help safeguard forests against increasing drought frequency.
5. *Synthesis*: Drought resistance and resilience of tree communities depend on tree diversity and mycorrhizal association types. Mixing tree species with diverse mycorrhizal types holds promise for forest restoration in the face of climate change.

## Introduction

Global warming leads to an increased likelihood of severe and consecutive droughts (Hari et al., 2020; Spinoni et al., 2018). Forest ecosystems, in particular, face threats from these droughts, given the long generation time and slow growth of trees compared to other plants (Brodribb et al., 2020). This is of great concern because, in addition to preserving biodiversity conservation and other ecosystem services, forests play a crucial role as carbon sinks, thereby contributing to the mitigation of present and future climate change (Anderegg et al., 2020).

In 2018, Northern and Central Europe experienced an extraordinary, compound drought, characterized by both, insufficient precipitation and heatwaves (Zscheischler et al., 2020; Zscheischler & Fischer, 2020). The exceptionally dry soil conditions persisted in 2019 and in vast areas of Central Europe, they lasted even through the year of 2020 (Rakovec et al., 2022). The occurrence of these three consecutive drought years from 2018 to 2020 (hereafter referred to as the ‘2018-2020 drought’) marks an unprecedented drought situation in Central Europe – at least within the last 250 years (Bastos et al., 2021; Hari et al., 2020; Rakovec et al., 2022). Such compound and consecutive droughts are substantially increasing in frequency (Hari et al., 2020; Markonis et al., 2021), and accumulating scientific evidence highlights their negative impacts on ecosystems, especially on forests ecosystems (Bastos et al., 2020, 2021; Forzieri et al., 2021; Gampe et al., 2021). Several studies reported widespread premature leaf senescence in 2018, unprecedented drought-induced tree mortality across various species throughout the region, and reductions in tree growth (Bose et al., 2020; Brun et al., 2020; Buras et al., 2020; Schnabel et al., 2022; Schuldt et al., 2020). Tree stress responses were found to be even more pronounced in 2019 than in 2018, indicating that consecutive and compound drought years represent a novel stressor for forests (Schnabel et al., 2022). Lags in physiological recovery, i.e. drought legacy effects, can be caused by hydraulic damage (Anderegg et al., 2018; Kannenberg et al., 2019), carbon depletion, or shifts in carbon allocation (e.g. towards rebuilding the canopy, growing roots or reproduction), which manifest in the reduction of radial stem growth.

For the quantification of drought impacts, the concept of resistance and resilience could be applied to forest communities to disentangle these different facets of ecosystem stability (Ingrisch & Bahn, 2018; Isbell et al., 2015; Lloret et al., 2011). Resistance is considered as the ability to persist and maintain functioning during a disturbance and can be quantified as the ratio between tree growth during drought and tree growth during the respective pre-drought period, characterized by ‘normal’ climate conditions. Resilience, which is defined as the capacity to reach pre-disturbance performance levels, is estimated as the ratio between post-drought growth and pre-drought growth (Lloret et al., 2011). Hence, assessing the differences in drought resistance and resilience among tree communities represents a crucial step toward understanding how drought impacts on forests could be mitigated by the choice of tree species and the design of climate-smart mixtures (Messier et al., 2022).

Biodiversity is known to stabilize ecosystem productivity over time and is therefore considered a key feature supporting the resistance and resilience of ecosystem functions to droughts (Cardinale et al., 2013; Isbell et al., 2015; Jourdan et al., 2020; Morin et al., 2014). In forest ecosystems, the influence of diversity on drought resistance and resilience is attributed to beneficial interactions among tree species such as resource partitioning (e.g. differential stomatal regulation strategies), facilitation (e.g. active hydraulic redistribution), or selection effects (e.g. competitive dominance of deep-rooted species) (Grossiord, 2020). In species-rich tree communities of the subtropics, the stabilizing effect of species richness has been experimentally confirmed and explained by the asynchronous growth dynamics of different tree species in response to variable climate conditions (Schnabel et al., 2021). However, whether this positive diversity effect persists when communities experience an unprecedented drought episode – such as the 2018–2020 drought – remains unknown. Indeed, there are indications that positive diversity effects observed under moderate drought stress may shift to negative effects due to competitive species interactions (Haberstroh & Werner, 2022). The evidence regarding the impact of tree diversity on forest growth during and after the drought remains inconsistent with positive, but also neutral and negative diversity effects on tree responses to drought being reported (Forrester et al., 2016; Gillerot et al., 2021; Grossiord, 2020; Grossiord et al., 2014; Jucker et al., 2014; Pardos et al., 2021).

One possible factor explaining these inconsistent results may be the type of mycorrhizal associations of the tree communities. Mycorrhizal fungi are known to assist plants in acquiring nutrients and water uptake in exchange for photosynthates (Bowles et al., 2018; Lehto & Zwiazek, 2011), as mycorrhizal hyphae reach soil water and nutrients that would be inaccessible to plant roots (Allen, 2007). Therefore, the type of mycorrhizal association could play an important role in drought effects on forests. A growing body of research suggests that the type of mycorrhizal association is a key driver for ecosystem-functioning relationships (Deng et al., 2023; Luo et al., 2023; Mao et al., 2023). There are two main groups of mycorrhizal association types that are formed between temperate tree species and fungi: ectomycorrhiza (EM) and arbuscular mycorrhiza (AM), differing in their morphology, physiology, and therefore soil nutrient uptake processes (Phillips et al., 2013; Tedersoo & Bahram, 2019). Ectomycorrhizal fungi develop a mantle of hyphae around plant root tips through which the nutrient exchange with their hosts occurs. Arbuscular mycorrhizal fungi are endophytic and exchange nutrients within the inner cortical cells of the plant host fine roots (Peterson & Massicotte, 2004). While AM fungi primarily provide their plant host with access to soil phosphorous in the upper mineral soil layer, EM fungi can mobilize both organic and mineral plant resources and typically thrive in organic soil horizons (Midgley & Phillips, 2014; Phillips et al., 2013; Read & Perez-Moreno, 2003; Rosling et al., 2016; Toju et al., 2016).

Although dual mycorrhization with AM and EM in plant roots seems common (Teste et al., 2020), one of the two mycorrhizal types dominates the association in most temperate tree species (Ferlian et al., 2021; Heklau et al., 2021). Due to the distinct lifestyles and foraging strategies of AM and EM fungi, it can be expected that the presence of both association types within a plant community could lead to higher resource partitioning among their associated plant hosts (Ferlian et al., 2018; Luo et al., 2018; Teste et al., 2020; Wagg et al., 2011b, 2011a). Especially during drought stress situations, in which tree communities lack water and nutrient supply, the potentially positive effect of mycorrhizal type richness could be more pronounced (Teste et al., 2020). However, evidence of the promising role of mycorrhizal associations for drought resistance and resilience of tree communities and especially studies using both mycorrhizal association types are lacking so far. For the determination of management actions for forests in future climatic conditions, it is crucial to understand the relevance of both, tree diversity and belowground mycorrhizal associations, for forest resistance and resilience to drought (Eisenhauer et al., 2022). Such insights could be best achieved with an experimental approach, which manipulates both factors while controlling for confounding environmental effects (Eisenhauer et al., 2022; Ferlian et al., 2018; Scherer-Lorenzen et al., 2005).

Here, we evaluated the growth resistance and resilience of tree communities varying in tree species richness to the 2018–2020 drought, using inventory data from a tree diversity experiment in Germany (MyDiv) that crosses tree species richness (monocultures, 2-species-mixtures, 4-species-mixtures) with mycorrhizal association types (AM only, EM only, AM + EM).

Specifically, we tested the following hypotheses:

> (H1) Growth resistance and resilience to the 2018-2020 drought increase with tree species richness.
>
> (H2) Communities containing tree species of both mycorrhizal association types (EM + AM) exhibit higher growth resistance and resilience to the 2018-2020 drought compared to tree communities with either arbuscular mycorrhizal (AM) associations or ectomycorrhizal (EM) associations alone.
>
> (H3) The relationship between growth resistance and resilience to the 2018-2020 drought and tree species richness is modulated by the mycorrhizal association type of the tree communities.

## Material and Methods

### Study site and experimental design

This study was conducted in the MyDiv Experiment, a tree diversity experiment located at the Bad Lauchstädt Experimental Research Station of the Helmholtz Centre for Environmental Research–UFZ, close to Halle (Saale) in Germany (51°23’ N, 11°53’ E; 118 m a.s.l.). The climate at the site is continental summer-dry with a mean annual precipitation sum of 484 mm and a mean temperature of 8.8°C (1896-2003). The soil at the site is a Haplic Chernozem developed from loess (Altermann et al., 2005) and was formerly used agriculturally until 2012 and as grassland until the establishment of the experiment in 2015. The study site was divided into two blocks based on the abiotic and biotic parameters measured before planting (Ferlian et al., 2018). The experiment consists of 80 plots, each 11 m x 11 m in size. In each plot, 140 two-year-old tree saplings were planted in a regular grid with a distance of 1 m between individuals. Tree species were selected from a species pool of ten common temperate deciduous tree species, with five identified in the literature as predominantly associating with EM fungi and five predominantly with AM fungi (Table S1). The experiment is designed with a tree species richness gradient ranging from monocultures to four-species mixture plots (1, 2, 4 species), crossed with the mycorrhizal type, i.e. communities with AM tree species only, EM tree species only, or both, AM and EM tree species (AM+EM). Although the mycorrhizal design was established only indirectly by the selection of tree species, a 2019 study (Ferlian et al., 2021) provided confirmation that EM and AM trees are predominantly colonized by EM and AM fungi, respectively.

### Identification of drought period

We define drought as a period with higher water deficits in comparison to normal conditions, i.e. long-term means of meteorological parameters (Schwarz et al., 2020). We used the Standardized Precipitation Evapotranspiration Index (SPEI; Vicente-Serrano et al., 2010) and soil moisture patterns for the determination of drought years at our site. With this approach, we could quantify drought severity at different time scales based on a commonly used and standardized index, while also considering the importance of local soil conditions for plant-available water (Schwarz et al., 2020).

SPEI series were calculated with the SPEI package (Beguería et al., 2014) in R (R Core Team, 2023) from monthly precipitation (mm) and potential evapotranspiration (mm) data derived from the weather station located closest to the study site and with continuous records (DWD Climate Data Center [CDC], Station Leipzig/Halle, ID 2932; Figure S2). The last 40 years (1982-2022) were used as a reference period. With the calculated SPEI values, we classified the years between 1982 and 2022 into normal (SPEI ≥ (-1) | ≤ (+1)), particularly dry (SPEI ≤ (-1)), or particularly wet years (SPEI ≥ (+1)) (Figure S2). The years 2016 and 2017 can be considered ‘norma’ years, as their SPEI values are between (+1) and (-1). The years 2018, 2019, and 2020 had the lowest SPEI values in the last 40 years, i.e. we identified them as three years of consecutive severe drought. The year 2021 can be considered a particularly wet year, according to its SPEI value higher than +1. On site soil moisture data since 2017 that were collected by loggers directly on the experimental plots confirmed this drought year identification based on the SPEI (Figure S2).

### Tree growth responses

#### Tree measurements

In annually repeated tree inventories from 2015-2021, individual tree stem diameters were measured 5 cm above the ground with a diameter tape (basal diameter, d0; cm) in all 80 plots of the experiment. To avoid potential edge effects, we only used an area of 6 x 6 m and 36 tree individuals per plot, resulting in 2880 trees in total.

#### Data cleaning and preparation

Measurement errors in tree inventory data with an annual resolution are common due to e.g. inconsistencies in the precise measurement position at the stem, inconsistencies in the selection of the measured main stem for trees with multiple stems, or by the breakage and re-growth of a new stem in the same year (Fichtner et al., 2018; Schnabel et al., 2019). We applied a correction procedure for 4.2% of the values in the dataset, i.e., we predicted tree basal diameter via constructing individual-based allometric models with the usage of the following additional variables (a) diameter at breast height (DBH) in case the tree was higher than 1.3 m, or (b) height in case the tree was lower than 1.3 m (see Supplementary Section III for more details).

Before analysis, we excluded tree individuals with incomplete measurement series over seven consecutive years (2015-2021) (e.g. due to mortality, wind breakage, and re-sprouting in the next year). Additionally, we removed all tree individuals of one plot of the experiment (monoculture of *Betula pendula*) due to its overall high damage and mortality caused by a storm event. This led to the exclusion of 272 tree individuals (9%) from the dataset.

With this approach there is a potential to overlook poorly performing trees, which could either restrict or enhance observed diversity effects. In general, to safeguard against any bias introduced by mortality, we calculated a mortality variable for each plot and year. This variable reflects the cumulative basal area lost as a percentage of the total basal area on the plot and was incorporated into all models as an additive factor. Importantly, subsequent analysis revealed that mortality did not emerge as a significant factor in any of the models.

#### Community productivity

Using d0, we calculated the tree basal area increment (BAI) for each year as: BAI_year_ = (π×(d0_year_/2)^2^) − (π×(d0_year-1_/2)^2^), where d0_year_ is the tree’s diameter in the respective year and d0_year-1_ its diameter in the previous year. With the individual BAI, we calculated community growth responses per year as the mean basal area increment [cm²] of all trees growing in the core area of one plot (max. 36 individuals) in the respective year.

#### Resistance and resilience

Tree community growth responses to drought were expressed as drought resistance and resilience (Lloret et al., 2011). We based the calculation of these indices on the tree community’s mean basal area increment (BAI) in the period before (2016-2017; “*pre”*), during (2018-2020; “*drought”*), and after (2021; *“post”*) the drought. It’s important to note that the duration of the pre-disturbance reference period markedly influences the resulting resistance value (Schwarz et al., 2020). However, due to the trees being planted in 2015, our options were constrained. As recommended by Schwarz et al. (2020), we compared our results for different reference periods (2016-2017 vs. 2017 only) and the results were found to be consistent. Therefore, we stayed with the more robust two-year reference period of 2016-2017. With 2022 also experiencing a severe drought, we could only assign 2021 as the post-disturbance year. Acknowledging that the timeframe of one year may not be optimal for expecting full recovery after a 3-year drought, we refer to this index as *early* resilience. We calculated resistance, the capacity to withstand drought (Lloret et al., 2011) as:

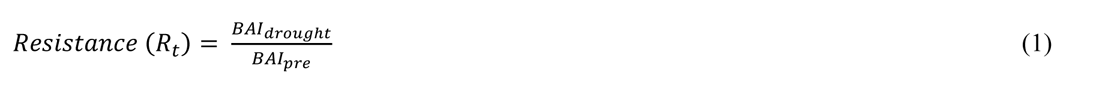

A resistance value below 1 indicates reduced growth during drought compared to the pre-drought period, while a value above 1 indicates increased growth in the drought period compared to the pre-drought period. However, given that we are examining very young tree individuals in the phase of exponential radial stem growth, we note that resistance values > 1 should be expected and a value around 1 represents an actual decline of growth.

We calculated early resilience, the ability to return to pre-drought conditions (Lloret et al., 2011), as:

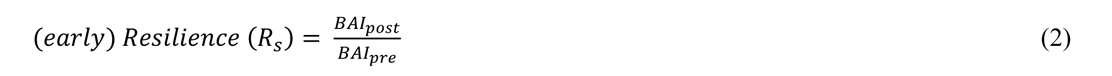

A resilience value of 1 indicates post-drought growth comparable to the pre-drought period and could result from either (a) high drought resistance or (b) low drought resistance coupled with strong recovery.

#### Community overyielding and species-specific overyielding

Using the individual BAI values per tree, we calculated the mean growth performance of the monocultures in the pre-drought, drought, and post-drought period for all ten tree species. Based on these values, we estimated the expected growth of mixed communities under the assumption that there would be no difference between the effect of inter-and intra-specific interactions (Forrester & Pretzsch, 2015). Since the trees were regularly planted with the same number of individuals per species in the mixtures (i.e. even mixing proportions per species), we used the mean over all monoculture performances per species as the “expected” community productivity in mixtures.

We quantified community-level over-or underyielding for each period as

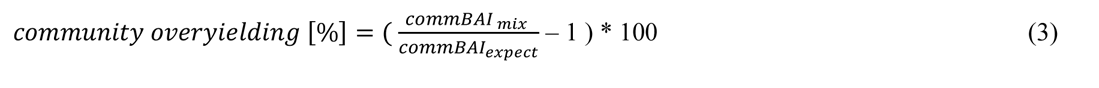

, where *commBAI_mix_* represents the actual community productivity in a two-or four-species mixture, and *commBAI_expect_* is the expected productivity of this community, calculated as the mean of all monoculture productivities of the species present in the respective community. When overyielding equals 0, the performance of the mixtures aligns with expectations based on the monocultures. Values above 1 represent improved performance in mixtures, indicating a benefit of mixing, while values below 1 indicate that the mixture performs worse than the corresponding monocultures.

Since a study on the same experimental site revealed that observed patterns of complementarity effects in tree growth were species-specific (Dietrich et al., 2022), we decided to gain some more deeper insights into the underlying mechanisms of the tree community behaviour during drought. Through a comparison of overyielding values among the different tree species, we aimed to disentangle the potentially contrasting mixing effects of individual species, which contribute to the diversity effect of the community.

To find out if a certain tree species benefits from growing in a mixture, in particular during drought, we calculated species-level overyielding as

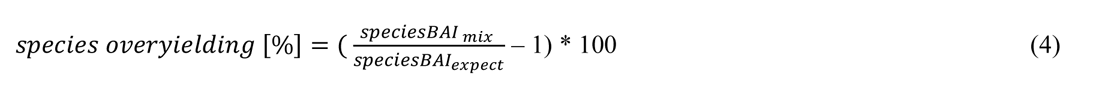

, using the species monoculture performance (as *speciesBAI_expect_*) and the species mean productivity in the respective community (as *speciesBAI_mix_*). A value above 1 indicates that a species benefits from growing in a certain mixture, compared to its performance in monoculture.

### Statistical analysis

To test our hypothesis that species richness and mycorrhizal association type shape tree community drought resistance and (early) resilience, we generated two linear mixed-effects models (LMM) for resistance and resilience, respectively. We modelled log-transformed resistance and resilience in response to the fixed effects of tree species richness (1, 2, 4; log-transformed), mycorrhizal type (as a factor: AM, AM+EM, EM), and their interaction. In addition, we included the mean tree basal area per community across years (tree size) and the accumulated lost plot basal area (mortality) as fixed effects to control for the effects of tree size and mortality and used the experimental block as a random effect.

Since we were not interested in examining the development of community productivity over the years and observed that absolute productivity was affected by individual species with high absolute growth rates (see Figure S6, Table S3), we focused on analysing relative responses. Consequently, we used an LMM to test whether the overyielding of the mixed communities depended on drought and mycorrhizal type using overyielding as the response variable, and drought period (pre, drought, post), mycorrhizal type (AM, AM+EM, EM), and their interaction as fixed effects. We further included species richness (as a factor: 2-, and 4-species) and mortality (as accumulated lost plot basal area) as additional fixed effects and controlled for repeated measurements by including the plot ID as a random effect.

We tested for overyielding in individual species and its dependence on period (pre, drought, post), through an LMM predicting species overyielding by interactive fixed effects of species identity (ten tree species), period (pre, drought, post) and tree species richness (2-, 4-species), with mortality (as accumulated lost plot basal area) as additional fixed effect and plot ID as a random factor. Further, to test if species differ significantly in their overyielding between the different drought periods, we used posthoc pairwise comparisons of drought periods within each species with the *contrast* function of the *emmeans* package (Lenth, 2023), corrected for multiple comparisons via Tukey’s Honestly Significant Difference (HSD) adjustment.

All analyses were conducted in R (version 4.3.1, R Core Team, 2023) using the packages *lme4* (Bates et al., 2015) for LMMs, *lmerTest* (Kuznetsova et al., 2017) for model selection via likelihood ratio tests, *performance* (Lüdecke et al., 2021) to check model assumptions, *emmeans* (Lenth, 2023) to extract model results and perform posthoc comparisons, and *ggplot2* for graphics (Wickham, 2016).

## Results

### Strong growth reductions during extreme drought

We found pronounced drought responses in terms of reduced tree growth during the 2018-2020 drought event. In the pre-drought period of 2016/2017, young trees exhibited an average growth (basal area increment) of 8.1 cm² (±2.7 cm²) (Table S4). However, over the three drought years, tree communities experienced an average growth reduction of 36.8 % compared to the pre-drought period (Figure 1). When comparing the individual drought years, the most extreme impact was observed in 2018, where tree communities exhibited the lowest average growth rate (4.8 cm² ±1.5 cm²), followed by 2019 (4.9 cm² ±1.7 cm²) and 2020 (5.7 cm² ±2.2 cm²). In the first post-drought year, 2021, the average growth rate slightly exceeded with 8.4 cm² (±2.1 cm²) the levels observed in the pre-drought period (Table S4). It is essential to note that young trees undergo a phase of exponential growth (Pretzsch, 2020), hence, rather than expecting constant growth, a notable increase in the growth rate would have been anticipated in 2018 and the subsequent years under ambient weather conditions.

**Figure 1:**
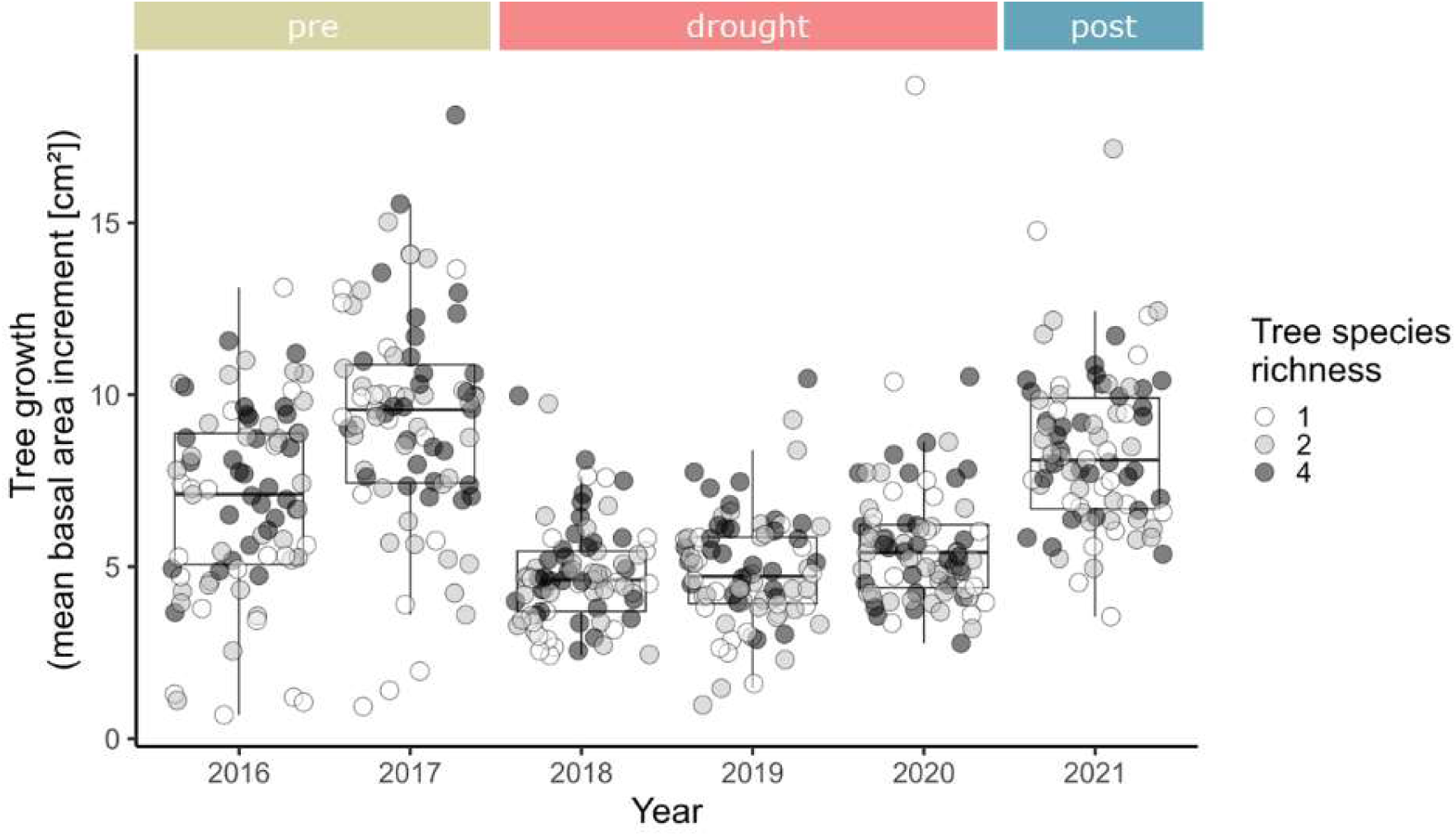
Mean annual growth of the tree communities per year. Shown is the mean tree basal area increment per plot over the years 2016 to 2021. The darkness of the grey colour shows the tree species richness level of the tree community. Boxplots for each year represent the interquartile range with the median indicated by the line inside the box.

Focusing on net diversity effects, we modelled their overyielding in relation to their respective monocultures across different periods of the drought event and with respect to the different mycorrhizal types of the plots. Our findings reveal that both the drought period and the mycorrhizal type significantly interacted as drivers of community overyielding (F_4, 114_ = 3.38, p<0.05; Table S5). The impact of whether the tree community consists of 2 or 4 different tree species on this relationship was hereby not significant. Before the drought, most diverse tree communities overyielded (Figure 2). However, during the drought period, the communities generally displayed reduced overyielding compared to the pre-drought period, approaching the performance levels of monocultures (indicated by the horizontal line, Figure 2). In the post-drought period, more communities showed underyielding than before or during the drought.

**Figure 2:**
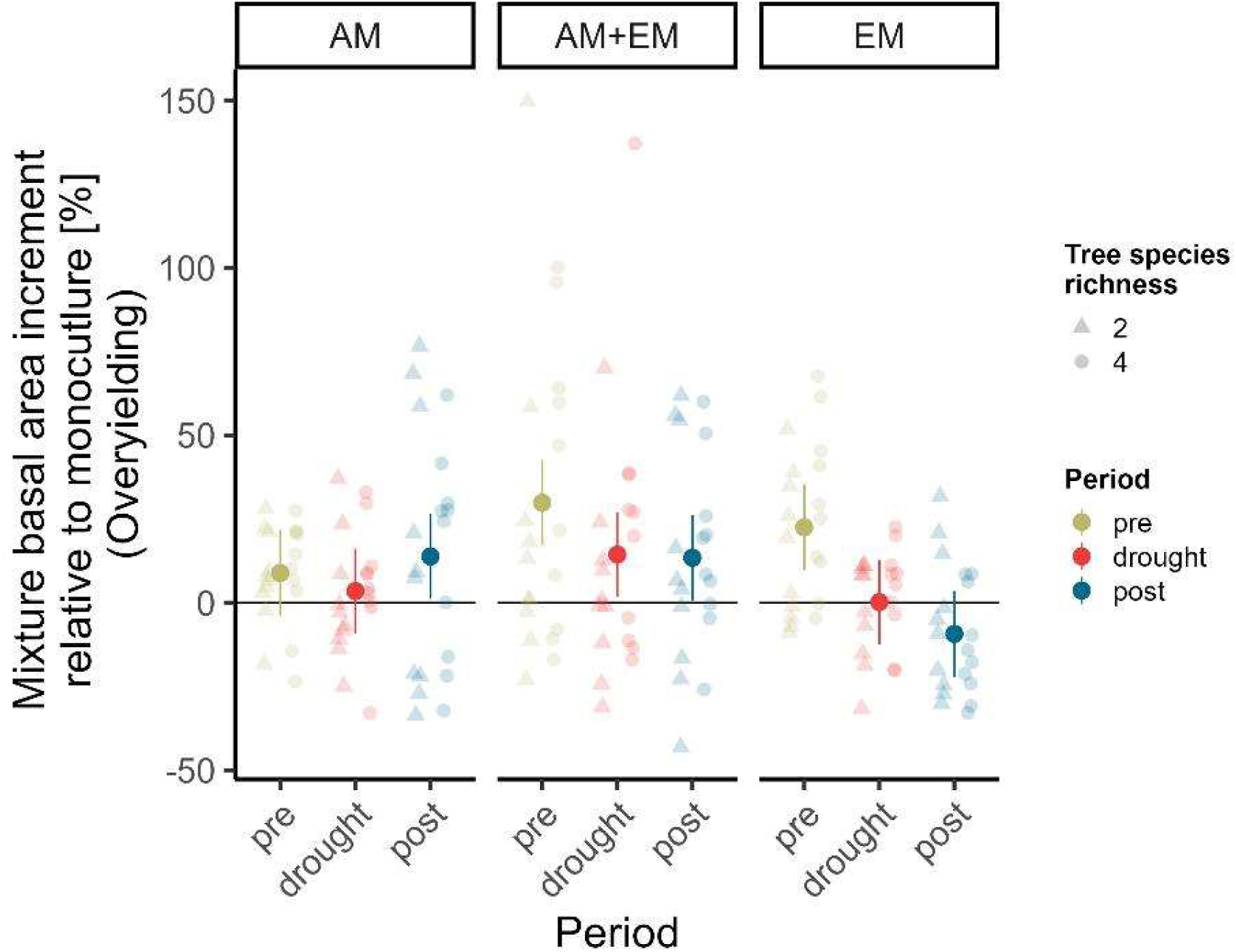
Overyielding of mixtures compared to monocultures. Shown is the basal area increment of mixed tree communities relative to their expected productivity based on the productivity of the constituent monocultures (black 0-line). Coloured points and error bars (95% CIs) show linear mixed-effects model fits that predict mean overyielding based on the examined period, tree mycorrhizal type, and species richness (see methods). The zero line represents the expected monoculture yield, i.e. values above this line indicate a mixture performance that is better than the respective monoculture, and values below this line indicate a mixture performance that is worse than the respective monoculture. The model explained 14% and 48% of the variation in overyielding through its fixed (marginal R^2^) and fixed and random effects (conditional R^2^). The different panels show the mycorrhizal community type (AM=arbuscular mycorrhiza; EM=ectomycorrhiza; AM+EM=both types).

Examining the different mycorrhizal types of the communities explained some of the shifts in overyielding patterns: in the pre-drought period, EM communities and communities with both mycorrhizal types, but not AM communities, clearly overyielded. We highlight that during the drought, only the communities with both mycorrhizal types (EM+AM) significantly outperformed their respective monocultures (Figure 2). In the post-drought period, the communities with AM trees and those with both mycorrhizal types predominantly displayed overyielding tendencies, whereas EM mixtures tended to underperform compared to their respective monocultures (Figure 2).

### Resistance and resilience modulated by species richness and mycorrhizal type

Overall, the tree communities had a drought resistance of 0.76 (± 0.5), i.e. they showed a decreased growth (R_t_ < 1) during the drought compared with the pre-drought period. Nevertheless, the same communities showed a mean drought resilience of 1.30 (± 1.2), i.e. they increased their growth after the drought compared to the period before the drought. For both responses, resistance and resilience, we found a high variability among the different communities (Figure 3). Overall, we found that mycorrhizal association type significantly shaped the relationship of tree species richness on both, the community’s drought resistance (F_2,70_ = 5.23, p = 0.0077, conditional R²=0.65) and resilience (F_2,70_ = 6.63, p = 0.0023, conditional R²=0.59)(Figure 3, Table S6). In contrast to our hypotheses, tree species richness did not consistently increase the drought resistance and resilience of tree communities (Table S6, Figure S7). Instead, tree species richness increased drought resistance and resilience only for AM communities and communities with both mycorrhizal association types, but decreased drought resistance and resilience for EM communities.

**Figure 3:**
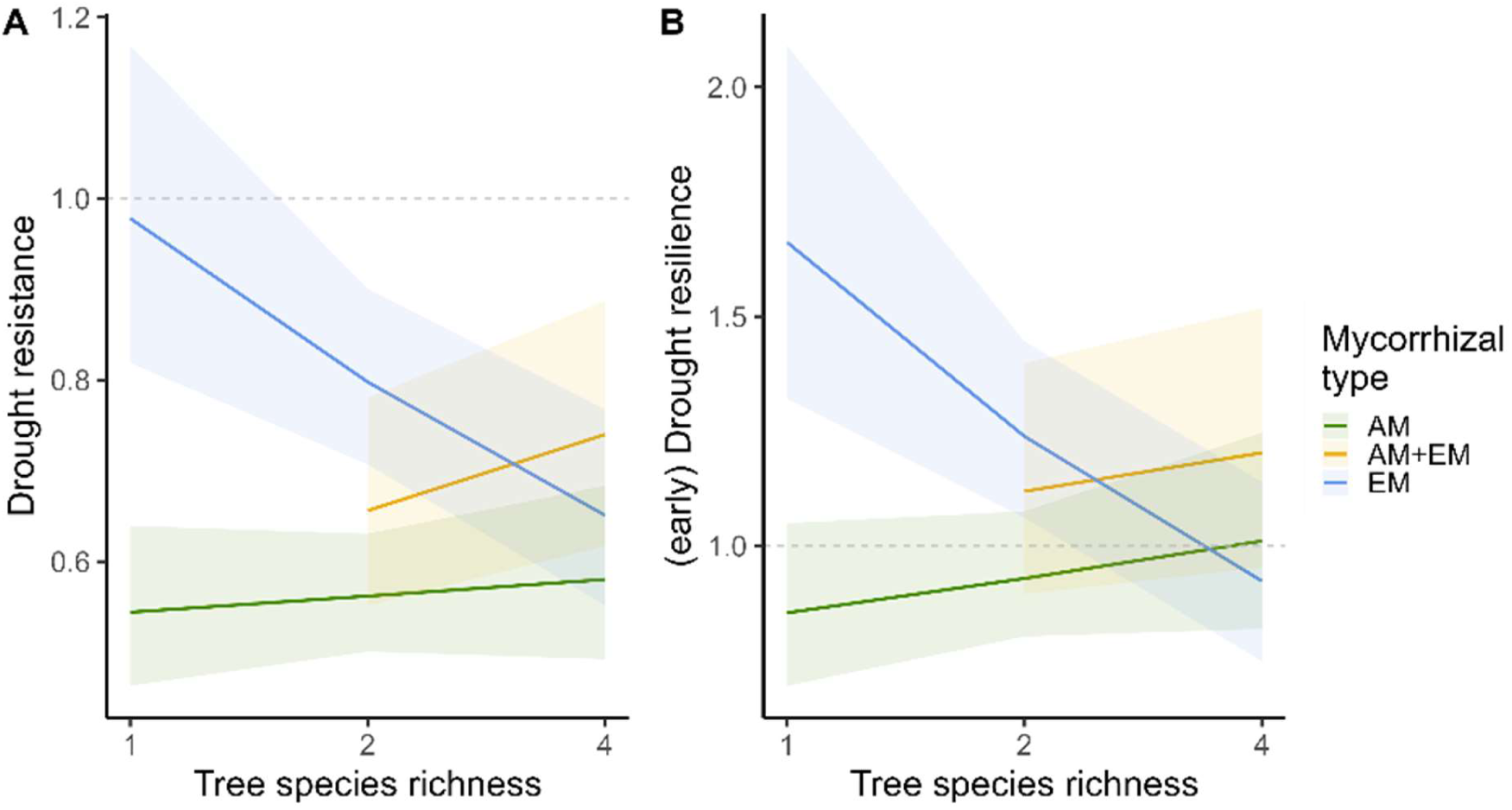
Tree community’s (A) drought resistance and (B) early resilience as a function of tree species richness and mycorrhizal association type. Colours refer to the plot’s mycorrhizal association type. Solid lines show significant (p≤0.05) linear mixed-effects model fits and shaded areas show confidence intervals of 95 % certainty. Dashed horizontal line at the intercept of y=1 as visual support for interpretation: values on this horizontal line are communities that grew as much during drought than before (resistance, A) or that grew as much after drought than before (resilience, B), values above the line stand for very high resistance and resilience, i.e. communities grew even more during drought than before (resistance, A) or more after drought than before (resilience, B).

In contrast to our hypothesis (H2), EM communities and not communities with mixed mycorrhizal types showed the highest drought resistance and resilience (Figure S7). Within the group of the EM communities, monocultures were the most resistant and resilient. However, focusing on tree species mixtures of the experiment, we found that while the mixed mycorrhizal communities showed an intermediate response between EM and AM communities in the 2-species mixtures, they surpassed both other groups in the 4-species mixtures (Figure 3). The relationship of drought resistance and resilience with tree species richness and mycorrhizal type was found to be additionally significantly dependent on the mean size of the trees in the community, with slightly higher resistance and resilience in small-sized trees (Table S6, Figure S8).

### Tree species identity determines the diversity benefit

We found that the benefit of diversity was strongly species-specific and depended on the drought period (Figure 4). Our model revealed that the interaction between species and drought period significantly predicted species overyielding in mixtures (F_18,404_ = 6.77, p<0.001, conditional R²=0.84). The degree to which a species benefited from the presence of other tree species in the community or not, was modulated by drought stress. Species such as *A. pseudoplatanus, P. avium, or B. pendula,* which already benefited from diversity under normal climatic conditions, experienced even greater advantages during drought. Conversely, species that did not benefit from diversity under normal climatic conditions, such as *A. hippocastanum, F. sylvatica, or Q. petraea,* tended to be even more negatively affected by diversity during drought conditions (Figure 4). These trends were more pronounced in both directions at the highest level of species richness for most species. Still, the lack of a significant modulation by tree species richness in the model implies that the observed effects remained relatively unaffected by the type of mixture (two-or four-species mixtures) (Table S8, Table S9).

**Figure 4:**
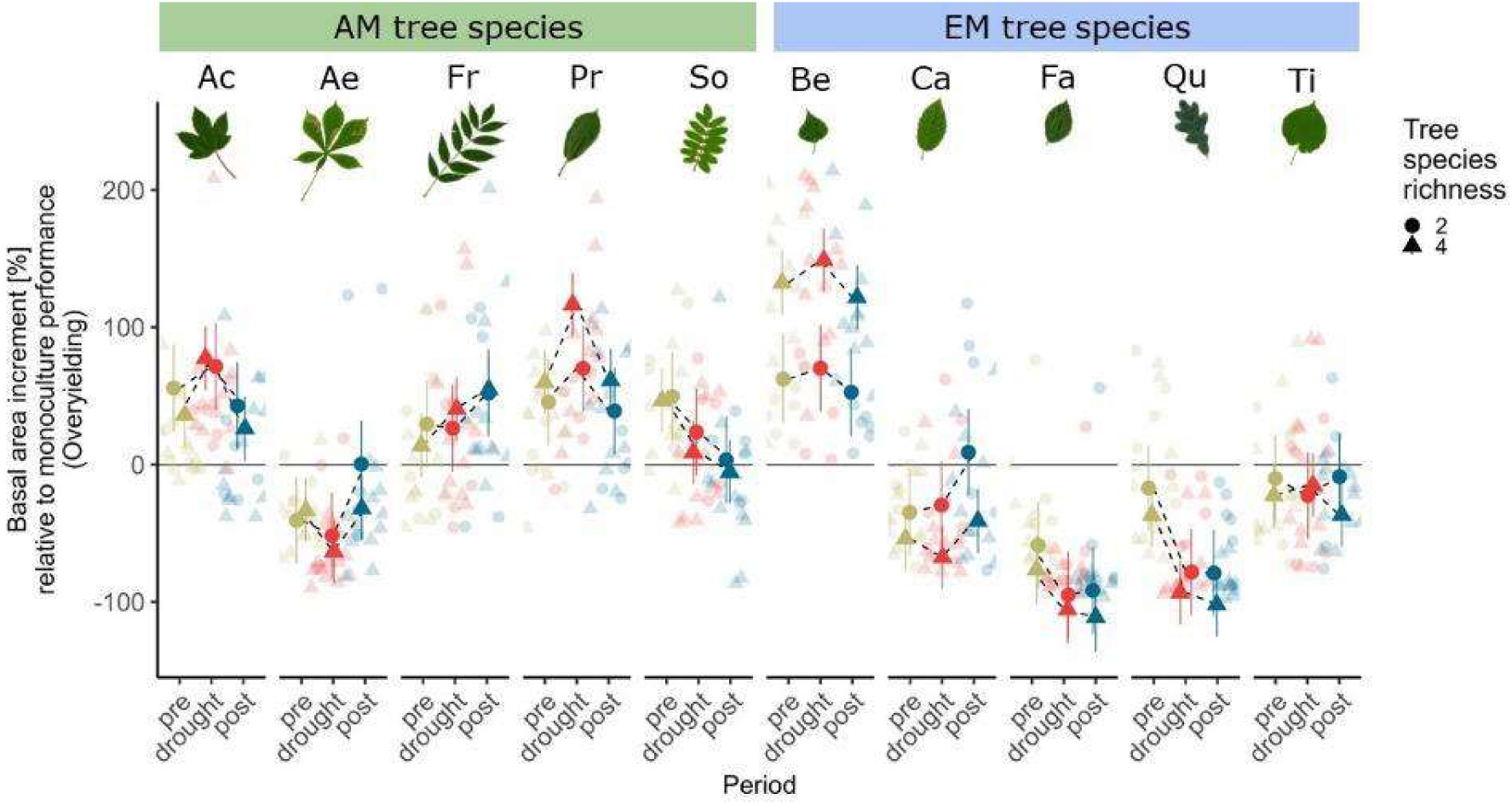
Overyielding of individual species in mixtures compared to monocultures. Each data point shows the basal area increment of a species on a mixture plot relative to its performance in monoculture (black 0-line). Colored points and error bars (95% Cis) show linear mixed-effects model fits that predict mean overyielding based on the examined period, tree species, and tree species richness (see methods). The zero line represents the expected monoculture yield, i.e. values above this line indicate a species performance in mixture that is better than the respective monoculture, and values below this line indicate a performance that is worse than the respective monoculture. The model explained 65 % and 84 % of the variation in overyielding through its fixed (marginal R²) and fixed and random effects (conditional R²). The different panels show which species is assigned to which mycorrhizal type. Species: Ac = Acer pseudoplatanus, Ae = Aesculus hippocastanum, Fr = Fraxinus excelsior, Pr = Prunus avium, So = Sorbus aucuparia, Be = Betula pendula, Ca = Carpinus betulus, Fa = Fagus sylvatica, Qu = Quercus petraea, Ti = Tilia platyphyllos.

## Discussion

### Strong growth reductions during extreme drought

We found strong declines in growth during the 2018-2020 drought event for the majority of the investigated tree communities (Figure 1). Other studies that assessed the impact of the 2018-2020 drought on forests have consistently reported signs of tree drought stress, reduced growth and increased tree mortality, both during and after the drought period (Buras et al., 2020; Obladen et al., 2021; Pohl et al., 2023; Schnabel et al., 2022; Schuldt et al., 2020; Senf et al., 2020). While our reported growth reduction by 36.8 % lines up with the finding of an earlier study by Thom et al., 2023, that reported a 41.3% reduction during the 2018-2020 drought, it should be kept in mind that our reductions are biologically even higher, because we expect an ontogenetic increase(Pretzsch, 2020). While a generally favourable nutrient supply can alleviate drought impacts on trees (Schmied et al., 2023), our initial anticipation of less pronounced growth reductions, based on the consideration of the soil characteristics at the site (Altermann et al., 2005) – particularly its high fertility and favourable water relations (see Methods) – did not align with the observed outcomes.

In terms of meteorological severity, out of the three drought years, 2018 stood out as the most extreme, followed by 2019 and 2020 (Hari et al., 2020; Rakovec et al., 2022) (Figure S1-S3). The growth responses of the tree communities at our site seemed to closely mirror the meteorological drought patterns, with the lowest growth rate in 2018, followed by 2019 and 2020. However, contrary findings from other studies point to 2019 as the year with the strongest growth reduction (Pohl et al., 2023; Salomón et al., 2022; Schnabel et al., 2022), attributed to the legacy effects of the 2018 drought (Anderegg et al., 2015; Kannenberg et al., 2020; Schnabel et al., 2022). Such drought legacies might be caused by diminished reserves of non-structural carbohydrates and altered carbon allocation, favouring the canopy and root system instead of radial stem growth (Hartmann & Trumbore, 2016; Brunner et al., 2015; Kannenberg et al., 2019). However, compared to most other studies, we studied young tree communities (planted in 2015; 5-6 years old when the drought hit), and the growth reduction we observed in 2019 may be interpreted as a strong reduction relative to the exponential growth trajectory expected for young trees (Pretzsch, 2020). Moreover, the less developed rooting system of young trees may have increased their sensitivity and led to earlier drought stress compared to trees in a mature stage, which may explain the pronounced reduction observed in 2018 (Franceschini & Schneider, 2014).

The year following the drought – 2021 – was a year with exceptionally high water supply (Figure S1, Figure S2). Nevertheless, we observed that tree growth remained lower in 2021 than in 2017 - the year immediately preceding the onset of the drought. Since the analysed trees experienced three consecutive extreme drought years, it is not surprising that they did not recover completely after a single year – whether this was despite, because of, or independently unaffected by their young age, we cannot be clarified with our data.

### Tree diversity did not increase drought resistance and resilience *per se*

In contrast to our hypothesis, we did not find that tree species richness *per se* can buffer the negative impacts of the extreme drought in 2018-2020. Neither drought resistance nor drought resilience of the investigated tree communities increased consistently with rising levels of tree species richness. Former studies provided mixed results for diversity effects on tree responses to drought (Dănescu et al., 2018; Gillerot et al., 2021; Grossiord, 2020; Jourdan et al., 2020; Pardos et al., 2021). The contrasting results on the role of tree diversity during drought may be explained by different drought-tolerance strategies of the admixed species (Schnabel et al., 2024) and the intensity of the examined drought event, as diversity effects on tree responses to drought may shift from positive to negative under severe drought (Haberstroh & Werner, 2022). Indeed, our expectation that the trees in mixtures would act complementarily in their resource use (especially water use) and consequently experience less drought stress and growth reductions compared to monocultures on the same site could not be confirmed. One possible explanation could be that during extreme water scarcity – such as during the 2018-2020 drought – even trees with complementary resource-use strategies compete for water resources (Haberstroh & Werner, 2022). A potentially enhanced competition level in mixtures during extreme drought is also supported by or species-specific analysis we present below. Moreover, it is important to consider that the higher productivity of mixed communities – which was also found for the mixtures of this study (Dietrich et al., 2022) - may increase the demand for water (Ammer, 2019). Mixtures could become more vulnerable to drought, unless mitigated by improved water supply via facilitated uptake (Forrester, 2015). Under extremely dry conditions, such as during the 2018-2020 drought, facilitation mechanisms, such as belowground niche differentiation, may no longer have been able to sustain tree water supply due to excessively dried-out soils. In these situations, the potentially advantageous larger root system of larger trees (Hui et al., 2014) within a mixture, would not confer benefits either. Our results reveal a similar response pattern for drought resistance as for early drought resilience (Figure 3). While we cannot confirm any positive effect of tree diversity on resilience (Anderegg et al., 2018), our ability to draw comprehensive conclusions on drought resilience is limited by the fact that our investigation only spans one year following the three consecutive drought years.

While various studies emphasize the influence of tree species composition on drought responses in mixed versus monospecific stands (Dănescu et al., 2018; Gillerot et al., 2021; Jourdan et al., 2020; Pardos et al., 2021), the distinctive strength of our study emerges from being the first to examine impacts of the 2018-2020 drought under the controlled conditions of a planted tree diversity experiment with various combinations of tree species exposed to the same abiotic conditions. Capitalizing on this setup, we revealed that the observed drought effects were not only driven by tree species richness but rather depended on the mycorrhizal association type of the examined tree communities.

### Mycorrhizal types modulate drought responses

In line with our expectations, we found different responses to drought in AM and EM tree communities, but with higher resistance and resilience for EM tree communities. There is evidence that the mycorrhizal association types differ in their nutrient economy: AM fungi rely on inorganic nutrient resources, and EM fungi have the ability to decompose organic matter (Averill et al., 2019; Deng et al., 2023; Liese et al., 2018; Phillips et al., 2013; Tedersoo & Bahram, 2019; Zhang et al., 2018). However, given that the study site has relatively high soil nutrient availability due to its former agricultural use (Ferlian et al., 2018), the differences between AM and EM in acquiring nitrogen and phosphorus, may not be decisive in this context. In particular, during drought, it might be more advantageous for the tree communities to have an enhanced water supply through their fungal partners than an improved nutrient supply. Both mycorrhizal fungi have mechanisms to maintain host vitality under drought, such as the induction of host aquaporin expression, regulating water uptake (Allen, 2007; Lehto & Zwiazek, 2011; Mohan et al., 2014; Tedersoo & Bahram, 2019; Xu & Zwiazek, 2020). The question of whether EM or AM associations offer greater drought resistance and resilience to their host trees remains uncertain, given the contrasting results reported so far and the lack of studies that compare both mycorrhizal types (Mohan et al., 2014; Querejeta et al., 2009). Some studies would support the benefits of AM associations under drought, such as AM hyphae being able to endure highly negative water potentials and having a greater plasticity of hyphal production, which might support the existence of AM host plants in extremely water-limited systems (Querejeta et al., 2007; Tedersoo & Bahram, 2019; Vargas et al., 2010). However, our results show that tree communities with EM associations are more resistant and resilient at our site. We propose the following speculative explanations for this response: EM fungi are expected to transport soil water more efficiently due to their greater mycelium biomass and their ability to build vessel-like rhizomorphs (Allen, 2007). Since EM fungi form a Hartig net of hyphae surrounding the root cortex cells and a hyphal sheath, covering the root tips (Freschet et al., 2021), they offer their host superior physical protection against soil-borne pathogens compared to AM fungi (Tedersoo & Bahram, 2019). This enhanced protection might be particularly advantageous during periods of stress, such as drought. Accordingly, it was observed that limited soil water supply reduced stem biomass production stronger for AM than for EM trees in a mesocosm drought experiment (Liese et al. 2018). Furthermore, the drought treatment only reduced fine root biomass and mycorrhizal colonization rates in AM trees, not in EM trees. On the broader scale of ecoregions, a study by Luo et al. (2023) found EM-dominated forest communities to be more productive in those ecoregions, where mean annual precipitation was low. Even though the mechanisms remain unclear, these findings support ours, as EM tree communities showed the highest drought resistance and resilience at our site. Still, while the evidence of the importance of mycorrhizal types for forest ecosystem-functioning relationships is accumulating (Deng et al., 2023; Dietrich et al., 2022; Luo et al., 2023; Mao et al., 2023), we need more knowledge on how these relationships are influenced by drought.

Interestingly, within the EM tree communities in our study, monocultures demonstrated greater resistance and resilience than mixtures, while the opposite trend was observed for AM tree communities, i.e. higher resistance and resilience were associated with higher tree species diversity. This could be explained by potentially lower root protection by AM fungi colonization compared to EM fungi colonization, resulting in AM trees suffering more from the accumulation of antagonists near conspecifics compared to EM trees (Bennett et al., 2017; Jiang et al., 2020). In addition, EM fungi appear to face fewer challenges in extending their growth from one root to another compared to AM fungi (van der Heijden & Horton, 2009). Given the host-specificity of mycorrhizal fungi (Ferlian et al., 2021), this ability might lead to a higher benefit from mycorrhizal fungal networks for EM tree monocultures.

Other studies on the same experimental site could not find any general beneficial effect of mixing mycorrhizal types but rather the tendency of an additive effect of EM trees and AM trees, e.g. in terms of biomass production (Dietrich et al., 2022; Ferlian et al., 2018) or stand structural complexity (Ray et al., 2023). However, when looking at the joint effects of tree species diversity and mycorrhizal type in our study, we found a strong positive slope only among the mixed mycorrhizal communities, leading to the highest resistance and resilience of mixed mycorrhizal communities within the 4-species communities (Figure 3). This indicates that complementarity effects under drought are enhanced (Baert et al., 2018) not only when tree species per se but also associated mycorrhizal association strategies are mixed. Our findings regarding community overyielding highlight that communities with mixed mycorrhizal types outperformed monocultures before, during, and after the drought (Figure 2), representing a novel drought mitigation effect of diversity. The fact that the modulation of mycorrhizal types differs considerably between periods may also be a reason why other studies on the same site, such as Ray et al. (2023), found no consistent mycorrhizal effect on productivity over the entire time period. While there is not yet much literature to support our findings, recent studies point in the same direction and provide the first evidence that the variability in mixed mycorrhizal strategies can promote ecosystem functioning. For instance, the study by Luo et al. (2023) in forest plots in the U.S. reported that communities with mixed mycorrhizal strategies were more productive than communities where either EM or AM tree species were dominant. However, our study is the first to demonstrate the positive effects of mixed mycorrhizal strategies related to drought responses. If confirmed in subsequent studies, this positive effect of mycorrhizal type mixtures would be a novel drought mitigation effect in forests, in addition to the positive effect of tree diversity reported in some former studies (e.g. Fichtner et al., 2020; Schnabel et al., 2019).

### Tree species’ benefits and disadvantages of diversity get stronger during drought

Our results revealed that diversity effects on productivity were highly species-specific. For instance, species like *A. pseudoplatanus*, *P. avium,* and *B. pendula* clearly benefited when growing in a mixture compared to their monocultures. On the other hand, species such as *A. hippocastanum*, *F. sylvatica,* and *Q. petraea* thrived more in monocultures than in mixtures. Species that benefitted from diversity in terms of overyielding grew even better, while species that underyielded in mixtures grew even less during drought than before or after the drought. These trends indicate that drought intensified the competitive differences among dominant and subdominant tree species. Our results suggest enhanced competitive dynamics during extreme drought, supporting the assumption that the positive effects of diversity may turn negative beyond a threshold of drought stress (Baert et al., 2018; Haberstroh & Werner, 2022) and emphasizing the need to consider drought intensity when discussing biotic interactions. Studies on drought responses of single tree species depending on the diversity of the community or neighbourhood are still rare. Some frequently-studied species, such as *F. sylvatica*, were found to benefit from diversity in terms of growth during drought for adult forest trees (Mölder & Leuschner, 2014; Vannoppen et al., 2020), which contrasts with our findings. But this advantage does not persist for *F.sylvatica* when mixed with conifers (Leuschner, 2020; Thurm et al., 2016), leading to the conclusion that diversity benefits are largely dependent on neighbour identity and neighbour size (Leuschner, 2020). Others state that the tree growth responses are contrasting depending on the drought intensity and the tree species (Bottero et al., 2021). In general, it seems to be evident that the influence of competition on tree growth responses during drought does not occur in an unidirectional and universal way for all species (Castagneri et al., 2022; Gillerot et al., 2021). Our results show that highly productive pioneer species, such as *B. pendula*, *P. avium*, or *A. pseudoplatanus*, suffered during drought in general (Figure S9); however, they benefitted more from growing in the mixture than the slow-growing species, such as *F. sylvatica* or *Q. petraea*. This observation is consistent with other experimental studies that reported that tree species richness particularly supported the most drought-vulnerable species in a community, characterized by acquisitive and water-spending functional traits (Fichtner et al., 2020; Schnabel et al., 2024).

It is important to note that the species-specific growth strategies cannot be clearly attributed to the mycorrhizal type, since growth strategies of e.g. EM tree species were found to cover a broad spectrum from the highest productivity across all species (*B. pendula*) to the slowest growth rate across all species (*F. sylvatica*), instead of a uniform strategy in all EM species (Dietrich et al., 2022). Despite the efforts to minimize variations in functional traits other than mycorrhizal type in the design of the experiment (Ferlian et al., 2018), we cannot dismiss the possibility that the observed patterns are partly linked to stand development dynamics. Furthermore, we assume that, alongside species-specific growth strategies, trait-based mechanisms – particularly those regulating water use – may explain the responses we observed. Since water regulation strategies are complex, a perspective of the whole plant (Hartmann et al., 2021) and therefore multiple (hydraulic) traits should be used in future studies to shed light on the drivers of drought resistance and resilience in addition to the mycorrhizal strategies studied here. Here, we only examined drought stress in terms of tree growth and disregarded other effects such as health deterioration or even mortality. Although the existing tree mortality on the experimental plots (Table S2) did not affect our results, low growth resilience of single communities may be an indicator of future mortality (DeSoto et al., 2020). To gain a more complete image of the 2018-2020 drought consequences, future studies should incorporate additional drought stress indicators, such as carbon isotope ratios (Cherubini et al., 2021; Jucker et al., 2017). Moreover, a thorough examination at the neighbourhood level, where tree-tree interactions occur (Trogisch et al., 2021) may provide a more comprehensive explanation of complementarity and competitive species interactions in the context of drought.

## Conclusion

Our results showed that it is not tree species diversity *per se* that modulates drought responses, but it shapes, in interaction with the mycorrhizal types and their diversity, the drought resistance and resilience. Our findings highlight that among various tree species mixtures, only those with mixed mycorrhizal types consistently exhibited overyielding during the extreme 2018-2020 drought. Even though we cannot elucidate the mechanisms behind the benefit of mixed mycorrhizal types during drought in the present study, important consequences can still be drawn from our observations. Our results highlight the potential of mixtures comprising tree species with different mycorrhizal types for effective forest restoration strategies, particularly in the face of an increasing frequency of extreme drought events like the 2018-2020 drought.

Further, we found that the drought-mitigating effect of diversity is most pronounced for fast-growing species, which overall suffered most from drought, as indicated by their comparably low drought resistance. We observed that drought intensified competitive differences among tree species, resulting in winners and losers under these harsh environmental conditions. Nevertheless, our comprehension of the processes and consequences of drought on interactions within tree species mixtures is still in its early stages. Additional experimental evidence is required for predicting the vulnerability of trees in the face of climate extremes. We emphasize the need for: (i) a thorough examination at the neighbourhood level, (ii) detailed information on the drought-tolerance traits of individual tree species to characterize their physiological strategies, thereby providing a better explanation of species interactions under drought, and (iii) studies that include the belowground perspective (e.g. the physiological strategies of fungal partners and their activity during drought) to understand what actually happens to the mycorrhizal symbiosis itself when soil water is limited.

Although the transferability of our results to mature forests is limited, our emphasis on young tree plantations remains particularly pivotal in the context of ongoing reforestation initiatives. Our study stands among the pioneering efforts to examine the severe impacts of the 2018-2020 drought in an experimental setup, where tree species richness and mycorrhizal type were manipulated. Capitalizing on this setup, we could directly compare the influence of the mycorrhizal types on the tree communities’ drought responses under similar abiotic conditions. Though the topic of mycorrhiza during drought requires further investigations, our findings already imply that a mixture of mycorrhizal types within tree communities may be a promising strategy for safeguarding forests against increasingly frequent severe drought events under climate change.

## Acknowledgments

This study was supported by the International Research Training Group TreeDì jointly funded by the Deutsche Forschungsgemeinschaft (DFG, German Research Foundation) – 319936945/GRK2324 and the University of Chinese Academy of Science (UCAS). N.E. gratefully acknowledges the support of iDiv, which is funded by the German Research Foundation (DFG – FZT 118, 202548816), as well as by the DFG (Ei 862/29-1; Ei 862/31-1).

## Author contributions

C.W., F.S., and L.S. conceived the ideas and developed the concept of the study. N.E. and O.F. designed and established the experiment. J.Q. collected the data. L.S. and P.D. cleaned the data. L.S. analysed the data with support from F.S., R.R., and C.W. L.S. led the writing of the manuscript with support from

F.S. and C.W. All authors contributed critically to the drafts and gave final approval for publication.

## Data availability

Data will be made publicly available in the MyDiv data portal (https://mydivdata.idiv.de/) upon publication.

