## Supplementary File for "Forest growth resistance and resilience to the 2018-2020 drought depend on tree diversity and mycorrhizal type"

SECTION I: Experimental Setup.......................................................................................2

Table S1: Tree species in the MyDiv experiment

Table S2: Mortality within the different experimental communities in the MyDiv experiment

SECTION II: Drought Identification..................................................................................3

Figure S1: Precipitation and Temperature

Figure S2: SPEI Index

Figure S3: Soil moisture content

SECTION III: Data Cleaning Procedure & Data Preparation..........................................6

Figure S4: Example of the correction process for one tree individual

Figure S5: Histogram of the $R^2$ values of all the 5120 individual linear models

SECTION IV: Model Supporting Information..............................................................8

Figure S6: Community growth over the years

Table S3: Community Growth – Model Output

Table S4: Overview of growth in single years

Table S5: Community Overyielding – Model Output

Table S6: Resistance and Resilience – Model Output

Figure S7: Drought resistance and early drought resilience with only species richness as fixed factor or only mycorrhizal type as fixed factor

Figure S8: Correlation of drought resistance (A) and drought resilience (B) with the mean tree basal area

Table S7: Overview of drought resistance and drought resilience

Table S8: Species Overyielding – Model output.

Table S9: Species changing their overyielding pattern during drought.

Figure S9: Species resistance and resilience.

### SECTION I: Experimental Setup

**Table S1: Tree species in the MyDiv experiment.**

| Species abbreviation | Species | Family | Mycorrhizal type | Mean height in 2021 [m] | Mean growth rate (2016-2021) [cm²/year] | Mortality [%] (until 2021) |
|---|---|---|---|---|---|---|
| *Ac* | *Acer pseudoplatanus* L. | Sapindaceae | AM | 5.6 | 8.4 | 4.7 |
| *Ae* | *Aesculus hippocastanum* L. | Sapindaceae | AM | 3.2 | 4.5 | 2.9 |
| *Fr* | *Fraxinus excelsior* L. | Oleaceae | AM | 4.8 | 6.7 | 10.7 |
| *Pr* | *Prunus avium* (L.) L. | Rosaceae | AM | 5.7 | 9.7 | 1.4 |
| *So* | *Sorbus aucuparia* L. | Rosaceae | AM | 4.5 | 5.1 | 13.5 |
| *Be* | *Betula pendula* Roth | Betulaceae | EM | 6.0 | 10.9 | 21.9 |
| *Ca* | *Carpinus betulus* L. | Betulaceae | EM | 4.6 | 5.0 | 1.2 |
| *Fa* | *Fagus sylvatica* L. | Fagaceae | EM | 2.6 | 1.8 | 12.7 |
| *Qu* | *Quercus petraea* (Matt.) Liebl. | Fagaceae | EM | 2.8 | 2.1 | 12.1 |
| *Ti* | *Tilia plathyphyllos* Scop. | Malvaceae | EM | 4.5 | 8.0 | 2.0 |

**Table S2: Mortality within the different experimental communities in the MyDiv experiment.** Mortality in percent is calculated as: (i) the lost basal area of dead trees relative to the summed basal area of all alive trees per plot, accumulated over the years (referred to as: loss of basal area), and (ii) the number of dead tree individuals relative to the total amount of planted individuals at experiment's beginning (referred to as: missing trees). Non-accumulating values may result from replanting initiatives (until 2018) or regrowth of new shoots after wind breakage, but also - in case of loss of basal area – by increased biomass growth in surviving trees.

| | | *loss of basal area [%]* | *missing trees [%]* | *loss of basal area [%]* | *missing trees [%]* | *loss of basal area [%]* | *missing trees [%]* | *loss of basal area [%]* | *missing trees [%]* | *loss of basal area [%]* | *missing trees [%]* | *loss of basal area [%]* | *missing trees [%]* |
|---|---|---|---|---|---|---|---|---|---|---|---|---|---|
| **species richness** | **mycor-rhizal type** | 2016 | | 2017 | | 2018 | | 2019 | | 2020 | | 2021 | |
| 1 | AM | | 0.28 | 0.15 | 0.56 | 0.13 | - | 0.66 | 2.22 | 2.13 | 4.44 | 3.32 | 8.33 |
| 1 | EM | | 1.23 | 0.00 | - | 0.00 | - | 0.85 | 0.93 | 1.25 | 2.47 | 0.97 | 3.09 |
| 2 | AM | | - | 0.00 | - | 0.00 | - | 0.67 | 1.11 | 2.07 | 3.61 | 2.94 | 6.67 |
| 2 | AM+EM | | 0.83 | 0.29 | 0.83 | 0.24 | 0.28 | 2.55 | 6.11 | 3.75 | 8.06 | 3.27 | 9.17 |
| 2 | EM | | - | 0.06 | 0.28 | 0.04 | - | 3.35 | 3.33 | 5.12 | 4.72 | 4.82 | 5.83 |
| 4 | AM | | - | 0.00 | - | 0.01 | 0.28 | 1.28 | 2.50 | 2.28 | 5.00 | 3.08 | 6.67 |
| 4 | AM+EM | | - | 0.04 | 0.28 | 0.42 | 0.56 | 4.20 | 13.06 | 5.53 | 13.06 | 5.69 | 16.67 |
| 4 | EM | | - | 0.00 | - | 0.00 | - | 4.13 | 2.50 | 6.01 | 3.61 | 4.96 | 3.89 |

SECTION II: Drought Identification

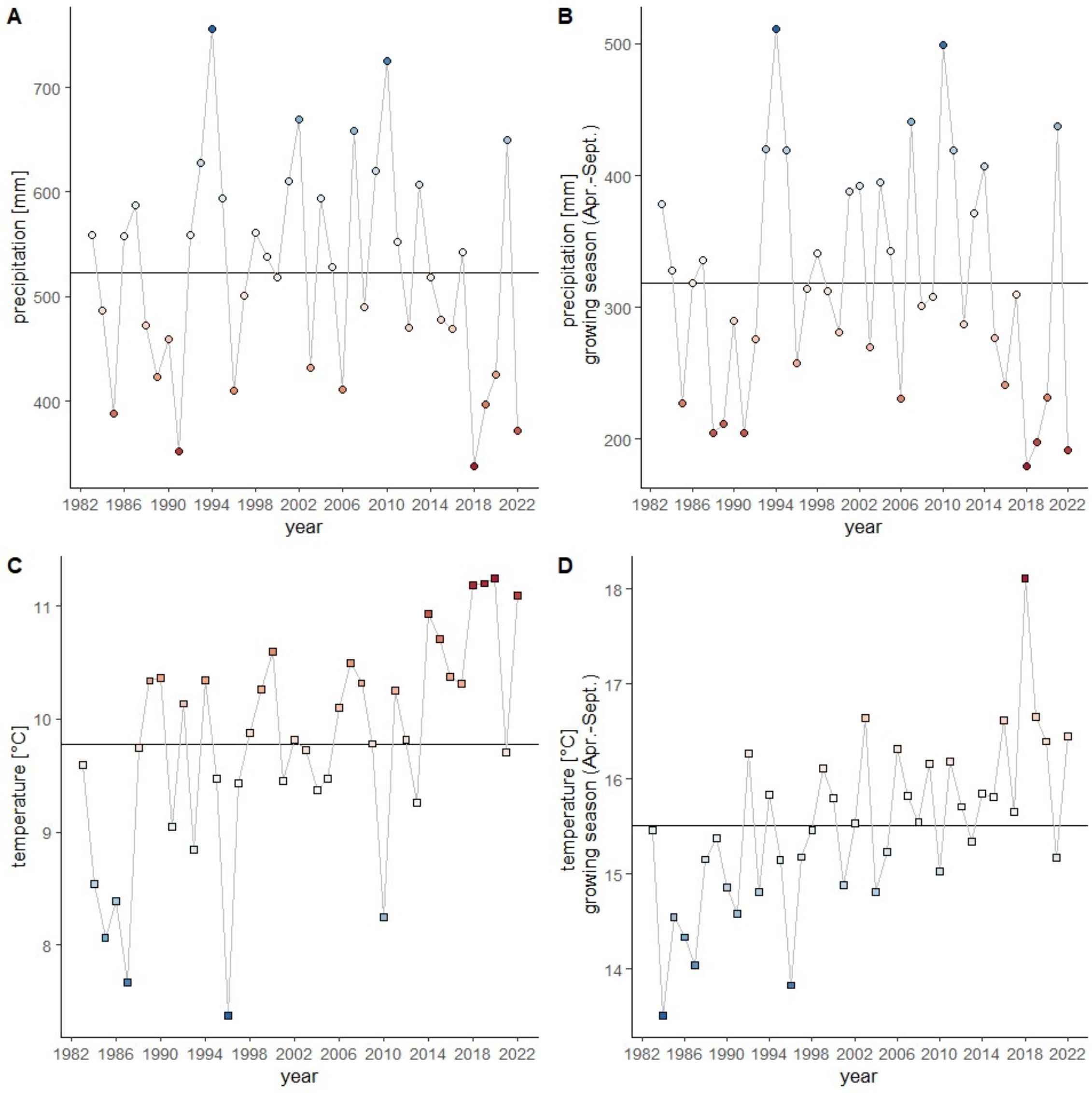

**Figure S1: Precipitation and Temperature.** Sum of precipitation and mean temperature in Leipzig/Halle from 1982 to 2022 for the whole year (A, C) and in the growing season (April-September; B, D). The horizontal line indicates the long-term mean over the shown period.

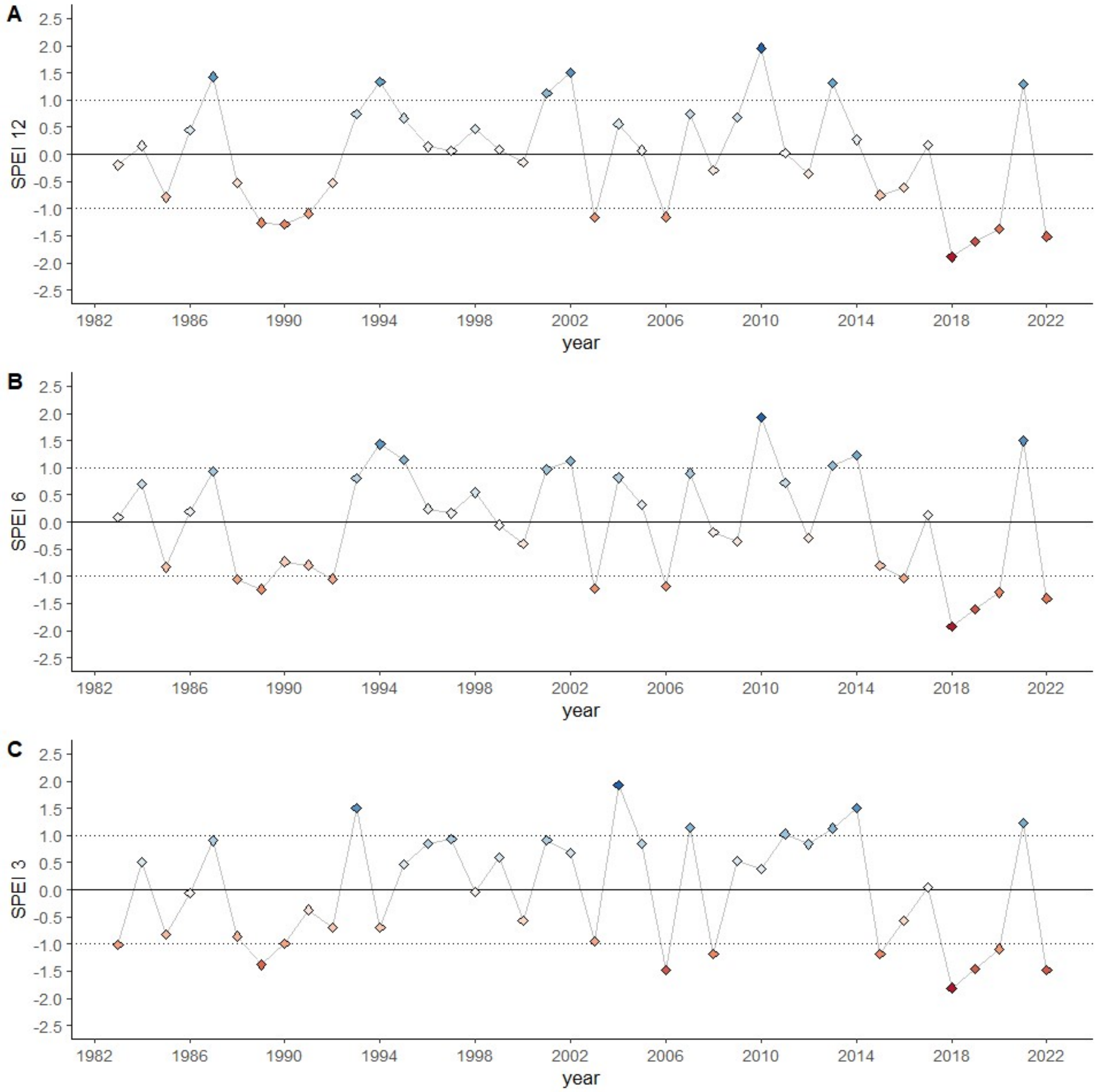

**Figure S2: Standardized Precipitation Evapotranspiration Index** (SPEI) for Leipzig/Halle from 1982 to 2022. Panels show three different time scales of SPEI calculation with (A) January-December (12 months) (B) April-September (6 months; growing season) and (C) May-July (3 months). SPEI values above and below the horizontal dotted lines (>(+1) or <(-1)) are considered as exceptionally wet and dry. The horizontal line at y= 0 represents the long-term mean.

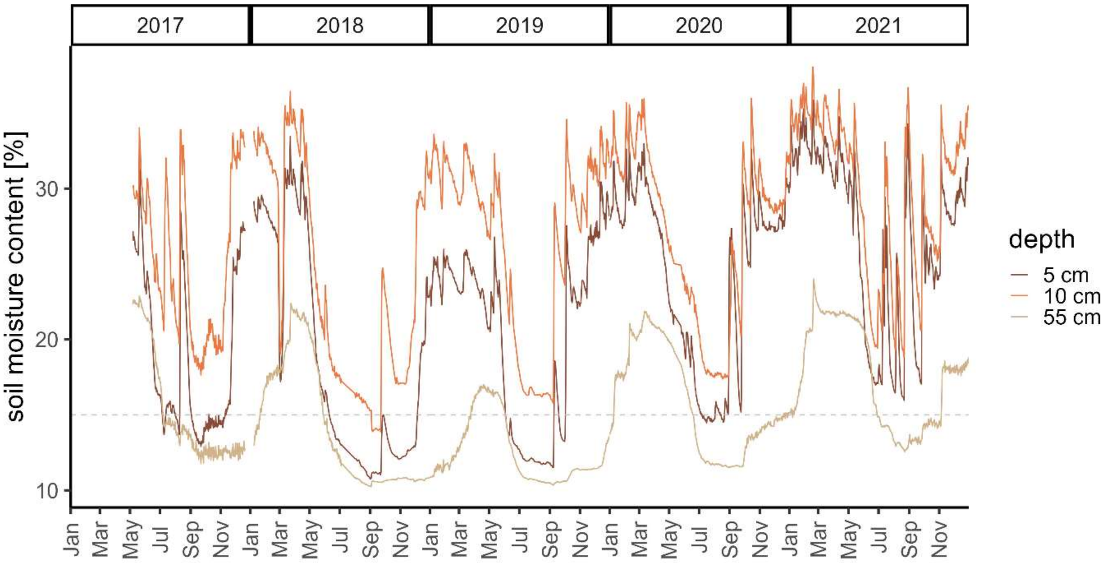

**Figure S3: Soil moisture content** [%] of the site since spring 2017. Daily mean values were derived from three different measurement loggers in the MyDiv experiment (at the center of plot 12, plot 57, and plot 77), measuring in 30 min intervals in three different soil depths (5 cm, 10 cm, 55 cm). The dashed horizontal line at 15 % shows the estimated permanent wilting point for the site (Altermann et al., 2005).

### SECTION III: Data Cleaning Procedure & Data Preparation

Measurement errors of the parameter diameter and height in tree inventory data with an annual resolution are quite common due to e.g. inconsistencies in the precise measurement position at the stem, inconsistencies in the selection of the measured main stem for multiple stem individuals, or by the breakage and re-growth of a new stem in the same year.

Initially, we computed an error distance for each measurement point by considering that the tree's diameter for a particular year should logically fall between the values for the diameter in the preceding and succeeding years. This assumption aligns with the unidirectional nature of tree growth. We calculated the error distance for a given year ($error\_distance_{year\_x}$) as the difference between the mean of diameters from the previous and subsequent years and the diameter of the tree in that specific year, divided by the diameter of the tree in that year:

$$error - distance_{year_x} = \left| \frac{mean(diameter_{year\ x-1}, diameter_{year\ x+1}) - diameter_{year_x}}{diameter_{year_x}} \right|$$

In cases where the error-distance value was higher than 0.5, we assumed a measurement error and identified this value as incorrect. Further, we categorized any diameter value as 'to be corrected' that showed negative increment compared to the preceding year. While acknowledging the occurrence of water-related contraction and expansion of wood and bark, we assume that actual shrinkage within a one-year timeframe, especially with measurements taken in usually well-watered winter conditions, is unlikely. This resulted in 1779 diameter values requiring correction, accounting for 4.96 % of the dataset. We corrected the selected values knowing about the usually tight relationship between the diameter at ground level, the diameter at breast height (DBH) and tree height. In pre-tests we found out that the relationships between the different measurement variables are not only highly species-specific, but also individual specific. Therefore, we build two linear regression models for each single tree individual (2 lm models x 5120 individuals) that predict the diameter at ground level by (1) the DBH and (2) the tree height (see **Figure S**).

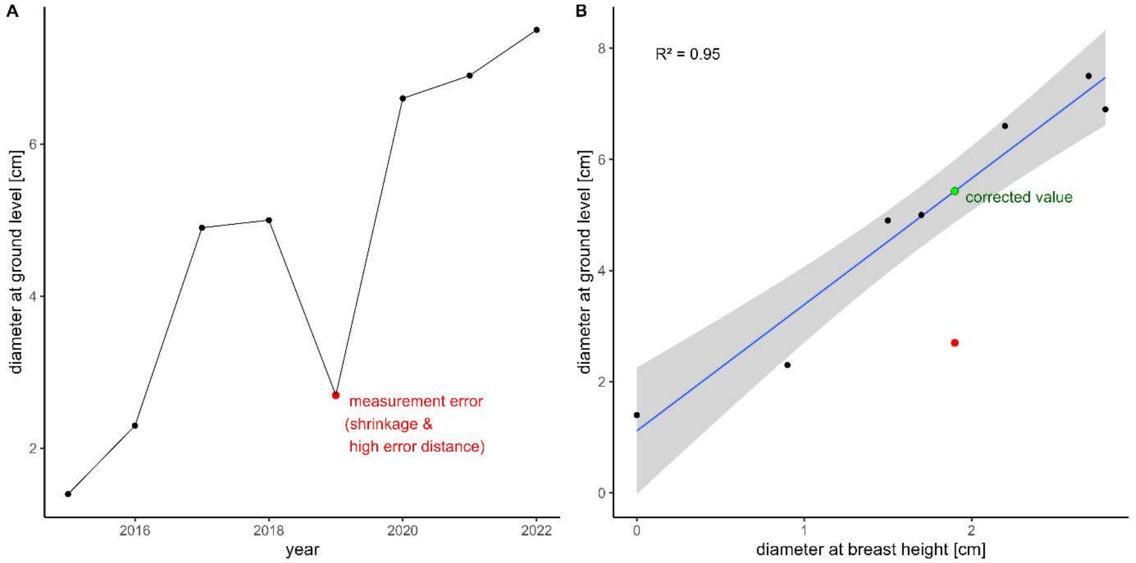

**Figure S4: Example of the correction process for one tree individual** of *A. pseudoplatanus* (Plot 36, Position H-05). (A) Growth in terms of diameter at ground level [cm] over the years with one datapoint identified as measurement error because of shrinkage in comparison to the previous year and a high error distance (datapoint marked in red). (B) Linear model of diameter at breast height [cm] predicting the diameter at ground level [cm]

with 7 datapoints of this tree individual. Original error value marked in red. New corrected value, predicted by the linear regression model is marked in green.

We took the prediction by the DBH model whenever the tree individual had a complete series of DBH values over all years (DBH could only be measured if the tree reached a minimum height of 1.3 m) and took in all other cases the predictions by the height model. Overall, the DBH models were more precise and succeeded in better model fits compared to the height models. This is why model fits were better for larger species that could more often be cleaned with the DBH model, because they were already higher than 1.3 m. However, some individual models could only explain very low variance proportion due to outlier values within their time series or missing data points and turned out to be not useful for the correction. We checked all the $R^2$ and whenever an individual model explained less than 50 % ($R^2$ below 0.5), we decided to keep the original value instead of correcting it (see **Figure S**).

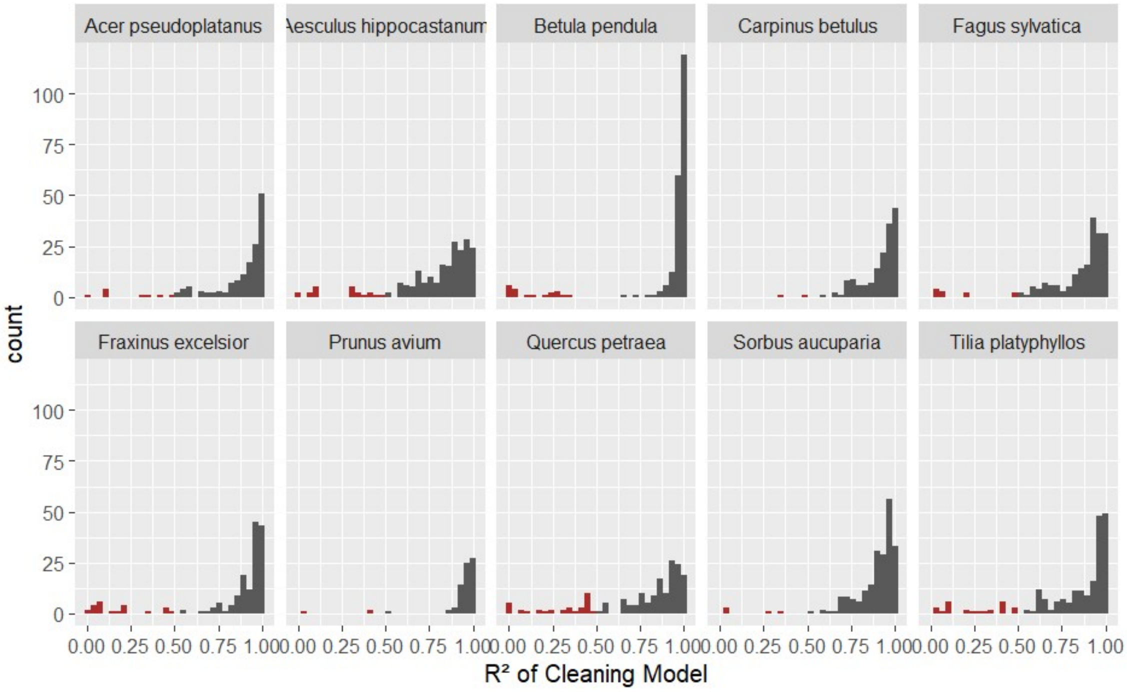

**Figure S5**: **Histogram of the $R^2$ values of all the 5120 individual linear models** (ground stem diameter predicted by DBH or by height, depending on whether DBH≠0) used for the cleaning process. All models with an $R^2<0.5$ (marked in red), were not used for the correction; instead the original value was kept. Different panels group the model fits for the 10 different tree species.

In the end we corrected 1596 diameter values (4.45 %) instead of the 1779 that we identified before as the ones that would have required correction.

To avoid potential biases in our data, we excluded tree individuals from the dataset under the following conditions: (a) those that were dead, even if they exhibited re-sprouting in the subsequent year, and (b) those that were once replanted, considering that until 2018, dead individuals were replanted. Additionally, we excluded trees located in the outermost row of the plot core area to adopt a more conservative approach regarding edge effects. This leads to considering up to 36 alive tree individuals within the center of the core plot area of 6 m x 6 m as a single tree community. The whole cleaning process resulted in a dataset of 2621 trees (instead of the 5120 planted ones) with complete growth series over all years.

### SECTION IV: Model Supporting Information

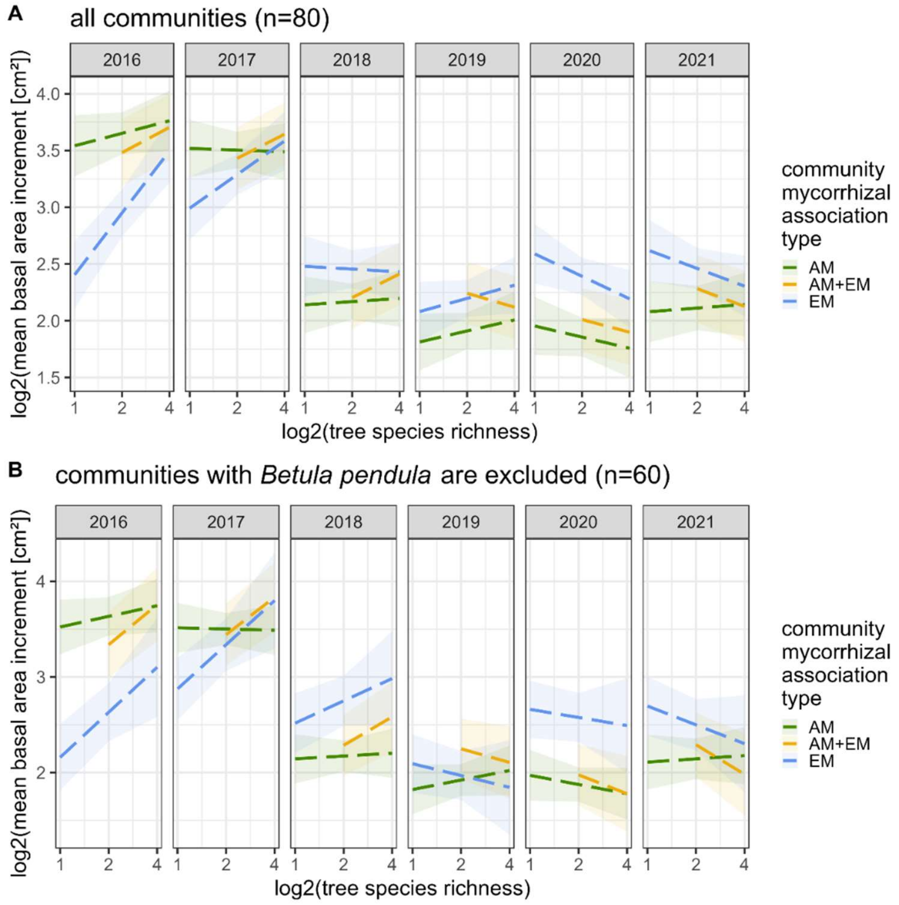

**Figure S6: Community growth over the years.** (A) Mean community basal area increments of trees on the central core area of the plots predicted by tree species richness and mycorrhizal type of the community. Linear regression lines show output of a linear mixed-effect model (LMM) that is only marginally significant (see Table S10) with 95 % confidence intervals. (B) The same model output than in A with the only difference that we excluded all communities with the presence of one very productive EM species, *B. pendula*, to see if the direction of effects are changing when loosing productivity in absolute terms. In 2018 and 2019 the regression line of EM communities is indeed changing the direction of the slope. Consequently, we decided that the model in A is overly sensitive to the contribution of single species in absolute terms.

**Table S3: Community Growth – Model Output:** Linear mixed-effect model (LMM) predicting mean growth of the trees within one central core area of a plot (log-transformed to meet model assumptions) with interactive fixed effects year (year: 2016-2021), mycorrhizal type of the plot (myc_type: AM, AM+EM, EM), log-transformed tree species richness of the plot (SR_log2: 0,1,2) and the additional fixed effect plot mean tree size (scaled). We used the plot id of the experiment as a random effect. Significant factors are printed in bold and asterisks additionally show the significance level in codes (p<0.001 '***', p< 0.01 '**', p<0.05 '*', p<0.1 '.')

| *predictors* | *Sum of Squares* | *Mean Square* | *Numerator DF* | *Denominator DF* | *F-value* | *p-value* | |
|---|---|---|---|---|---|---|---|
| *log(species richness)* | 0.50 | 0.50 | 1 | 437 | 2.61 | 0.1069 | |
| *mychorrizal type* | 0.04 | 0.02 | 2 | 437 | 0.10 | 0.9071 | |
| ***year*** | 14.17 | 2.83 | 5 | 437 | 14.71 | **0.0000** | *** |
| ***plot mean tree size*** | 36.39 | 36.39 | 1 | 437 | 188.89 | **0.0000** | *** |
| *log(species richness) : mycorrhizal type* | 0.30 | 0.15 | 2 | 437 | 0.78 | 0.4583 | |
| ***log(species richness) : year*** | 3.78 | 0.76 | 5 | 437 | 3.93 | **0.0017** | ** |
| ***mycorrhizal type: year*** | 14.70 | 1.47 | 10 | 437 | 7.63 | **0.0000** | *** |
| *log(species richness) : mycorrhizal type : year* | 3.50 | 0.35 | 10 | 437 | 1.82 | 0.0558 | . |
| *marginal $R^2$ / conditional $R^2$* | 0.597 / 0.597 | | | | | | |

**Table S4**: **Overview of growth in single years** per period and species richness. Tree growth as the mean of all individual tree basal area increment in cm² of alive trees growing on the core area of each plot. Means and standard deviations (sd) of all plots, or a subset of plots (monocultures, 2- and 4-species mixtures) in the different periods and years.

| | pre-drought | | | | drought | | | | | | post | |
|---|---|---|---|---|---|---|---|---|---|---|---|---|
| | *mean* | | *sd* | | *mean* | | | *sd* | | | *mean* | *sd* |
| *all plots* | **8.10** | | 2.65 | | **5.12** | | | 1.14 | | | **8.40** | 2.31 |
| | 2016 | | 2017 | | 2018 | | 2019 | | 2020 | | 2021 | |
| | *mean* | *sd* | *mean* | *sd* | *mean* | *sd* | *mean* | *sd* | *mean* | *sd* | *mean* | *sd* |
| *all plots* | **6.91** | 2.72 | **9.30** | 3.16 | **4.78** | 1.52 | **4.87** | 1.65 | **5.73** | 2.17 | **8.40** | 2.31 |
| *monocultures* | 5.57 | 3.35 | 8.59 | 4.07 | 4.33 | 1.57 | 4.10 | 1.23 | 6.35 | 3.47 | 8.05 | 2.72 |
| *2-species mixture* | 7.00 | 2.70 | 8.99 | 2.93 | 4.72 | 1.37 | 4.70 | 1.79 | 5.30 | 1.31 | 8.50 | 2.59 |
| *4-species mixture* | 7.66 | 1.98 | 10.06 | 2.63 | 5.13 | 1.62 | 5.53 | 1.51 | 5.75 | 1.72 | 8.53 | 1.72 |

**Table S5: Community Overyielding – Model Output:** Linear mixed-effect model (LMM) predicting overyielding (i.e. a plot's growth compared to its expected value calculated as the mean of the monocultures of the certain species composition) with fixed effects mycorrhizal type of the plot (AM; AM+EM; EM) and period of the drought (pre-drought; drought; post-drought) and their interaction, and the tree species richness (2species-mixture/4species-mixture) as well as the accumulated plot mortality (as basal area lost) (n=180). We used the plot ID of the experiment as a random effect. Significant factors are printed in bold and asterisks additionally show the significance level in codes (p<0.001 '***', p< 0.01 '**', p<0.05 '*', p<0.1 '.')

| *predictors* | *sum of squares* | *mean square* | *Numerator DF* | *Denominator DF* | *test statistic* | *p-value* | |
|---|---|---|---|---|---|---|---|
| ***period*** | 6962.40 | 3481.20 | 2 | 121.45 | 6.92 | **0.0014** | *** |
| *mycorrhizal type* | 2359.95 | 1179.98 | 2 | 56.09 | 2.35 | 0.1050 | |
| *species richness* | 574.42 | 574.42 | 1 | 56.11 | 1.14 | 0.2897 | |
| *mortality* | 53.00 | 53.00 | 1 | 168.12 | 0.11 | 0.7458 | |
| ***period : mycorrhizal type*** | 6797.29 | 1699.32 | 4 | 113.78 | 3.38 | **0.0118** | * |
| *marginal $R^2$ / conditional $R^2$* | 0.136 / 0.475 | | | | | | |

**Table S6: Resistance and Resilience – Model Output.** Linear mixed-effect models (LMM) predicting resistance and resilience with the fixed effects log-transformed tree species richness of the plot (SR_log2), mycorrhizal type of the plot (as factor: AM; AM+EM; EM) and period of the drought (as factor: pre-drought; drought; post-drought) and their interaction, and the mean tree basal area per plot over the years 2016-2021 (tree size) and the mortality during (year 2020) or after (year 2021) the drought (as accumulated lost plot basal area) as scaled additional fixed factors. We used the block id of the experiment as a random effect.

| ***log(resistance)*** | | | | | | | |
|---|---|---|---|---|---|---|---|
| *predictors* | Sum of Squares | Mean Square | Numerator DF | Denominator DF | F-value | p-value | |
| *log(species richness)* | 0.02 | 0.02 | 1 | 70.11 | 0.12 | 0.7342 | |
| *mycorrhizal type* | 3.95 | 1.97 | 2 | 70.02 | 13.96 | **0.0000** | *** |
| *tree size* | 7.04 | 7.04 | 1 | 70.93 | 49.80 | **0.0000** | *** |
| *mortality during drought* | 0.19 | 0.19 | 1 | 70.64 | 1.31 | 0.2560 | |
| *log(species richness) : mycorrhizal type* | 1.48 | 0.74 | 2 | 70.02 | 5.23 | **0.0077** | ** |
| *marginal $R^2$/ conditional $R^2$* | 0.636 / 0.647 | | | | | | |

| ***log(resilience)*** | | | | | | | |
|---|---|---|---|---|---|---|---|
| *predictors* | Sum of Squares | Mean Square | Numerator DF | Denominator DF | F-value | p-value | |
| *log(species richness)* | 0.12 | 0.12 | 1 | 70.18 | 0.50 | 0.4800 | |
| *mycorrhizal type* | 4.86 | 2.43 | 2 | 70.05 | 10.39 | **0.0001** | *** |
| *tree size* | 10.83 | 10.83 | 1 | 70.95 | 46.30 | **0.0000** | *** |
| *mortality after drought* | 0.28 | 0.28 | 1 | 70.99 | 1.19 | 0.2786 | |
| *log(species richness) : mycorrhizal type* | 3.10 | 1.55 | 2 | 70.03 | 6.63 | **0.0023** | ** |
| *marginal $R^2$/ conditional $R^2$* | 0.576 / 0.589 | | | | | | |

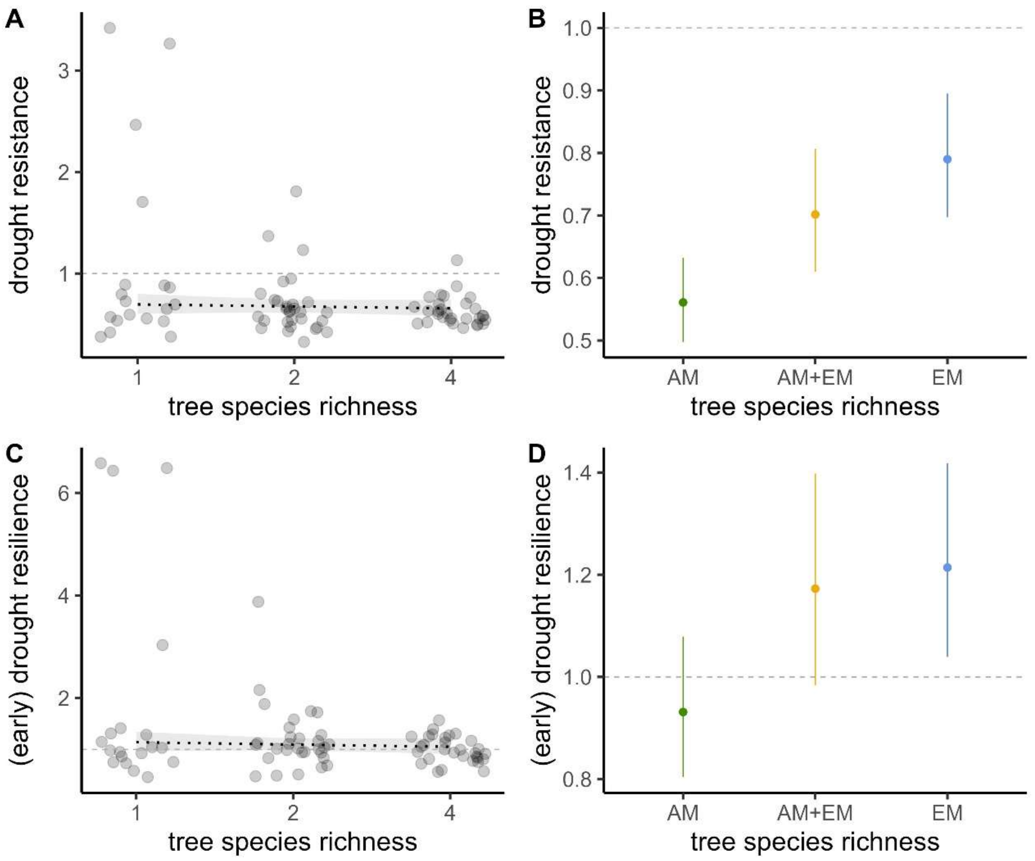

**Figure S7: Drought resistance (A, B) and early drought resilience (C, D) predicted by linear mixed effect models with only species richness as fixed factor (A, C) and only mycorrhizal type as fixed factor (B, D),** next to plot tree size and mortality as additional fixed factors and block of the experiment as a random factor. Response variables were log-transformed to meet model assumptions and back-transformed for the figures. Dashed lines indicate non-significant linear regression predictions.

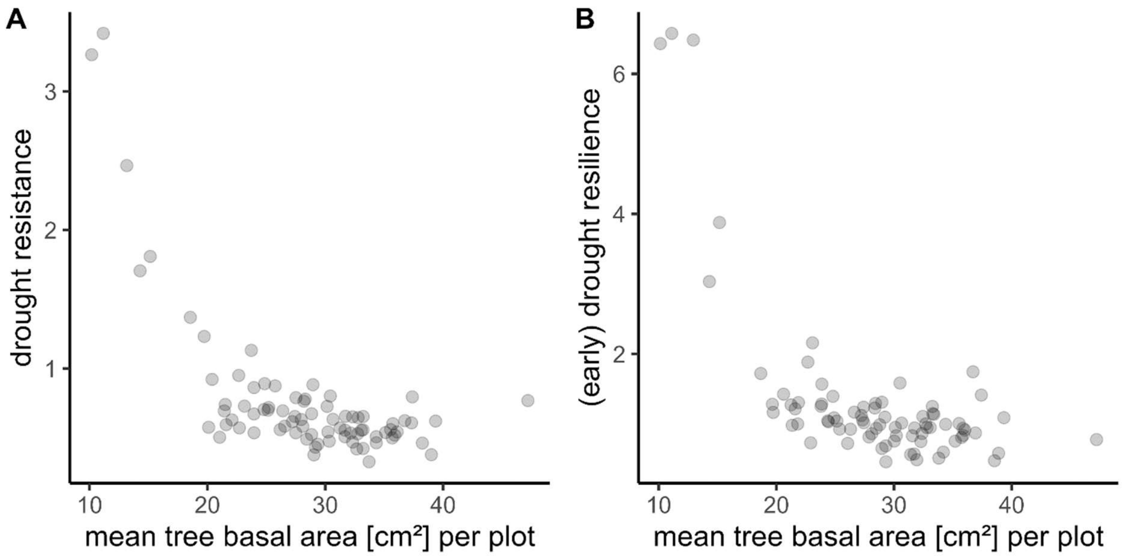

**Figure S8: Correlation of drought resistance (A) and drought resilience (B) with the mean tree basal area** [cm²] per plot (calculated as a mean value over the years 2016-2021)

**Table S7: Overview of drought resistance and drought resilience** per plot and experimental group (mean and standard deviation (sd). For resistance, single drought resistances for each drought year were calculated additionally, i.e. the drought year growth compared to the growth in the pre-drought level.

| *species richness* | | | ***1*** | | ***2*** | | | ***4*** | | | ***all plots*** |
|---|---|---|---|---|---|---|---|---|---|---|---|
| *mycorrhizal type* | | | ***AM*** | ***EM*** | ***AM*** | ***EM*** | ***AM+EM*** | ***AM*** | ***EM*** | ***AM+EM*** | |
| *N* | | | ***10*** | ***9*** | ***10*** | ***10*** | ***10*** | ***10*** | ***10*** | ***10*** | ***79*** |
| **resistance** | all drought years | mean | 0.57 | 1.62 | 0.51 | 0.88 | 0.68 | 0.54 | 0.74 | 0.61 | 0.76 |
| | | sd | 0.16 | 1.15 | 0.10 | 0.40 | 0.24 | 0.06 | 0.18 | 0.07 | 0.53 |
| | 2018 | mean | 0.53 | 1.30 | 0.55 | 0.91 | 0.61 | 0.52 | 0.64 | 0.60 | 0.70 |
| | | sd | 0.19 | 0.89 | 0.26 | 0.52 | 0.31 | 0.12 | 0.26 | 0.13 | 0.45 |
| | 2019 | mean | 0.52 | 1.35 | 0.46 | 0.76 | 0.69 | 0.56 | 0.72 | 0.60 | 0.70 |
| | | sd | 0.18 | 1.01 | 0.16 | 0.38 | 0.21 | 0.08 | 0.08 | 0.15 | 0.45 |
| | 2020 | mean | 0.66 | 2.22 | 0.53 | 0.99 | 0.72 | 0.54 | 0.85 | 0.64 | 0.88 |
| | | sd | 0.28 | 1.64 | 0.11 | 0.45 | 0.30 | 0.08 | 0.32 | 0.15 | 0.77 |
| **resilience** | all drought years | mean | 0.87 | 3.11 | 0.93 | 1.54 | 1.14 | 0.93 | 1.08 | 0.98 | 1.30 |
| | | sd | 0.28 | 2.62 | 0.48 | 0.89 | 0.32 | 0.31 | 0.26 | 0.17 | 1.15 |

**Table S8: Species Overyielding – Model output.** Linear mixed-effect model (LMM) predicting species overyielding with the fixed effects period of the drought (period: pre; drought; post), species (10 tree species) and tree species richness of the mixture (species richness: 2;4) and their three-way-interaction. Additional fixed effect is mortality as the accumulated lost plot basal area. We used the plot id of the experiment as a random effect. (significance level of α=0.05; p<0.001 '***', p< 0.01 '**', p<0.05 '*', p<0.1 '.')

| *predictors* | *Sum of Squares* | *Mean Square* | *Numerator DF* | *Denomi-nator DF* | *F -value* | *p-value* | |
|---|---|---|---|---|---|---|---|
| *period* | 5490.39 | 2745.20 | 2 | 420.12 | 2.52 | 0.0815 | * |
| ***species*** | 955104.13 | 106122.68 | 9 | 456.96 | 97.49 | **0.0000** | *** |
| *species richness* | 18.43 | 18.43 | 1 | 46.04 | 0.02 | 0.8970 | |
| *mortality* | 19.05 | 19.05 | 1 | 468.97 | 0.02 | 0.8948 | |
| ***period : species*** | 132691.90 | 7371.77 | 18 | 403.72 | 6.77 | **0.0000** | *** |
| *period : species richness* | 4238.89 | 2119.45 | 2 | 403.90 | 1.95 | 0.1440 | |
| ***species : species richness*** | 67818.21 | 7535.36 | 9 | 456.97 | 6.92 | **0.0000** | *** |
| *marginal $R^2$/ conditional $R^2$* | 0.653 / 0.835 | | | | | | |

**Table S9: Species changing their overyielding pattern during drought.** Post-hoc tests as pairwise overyielding contrasts of the drought periods within each species, adjusted p-values by tukey correction method for multiple comparisons. For information on underlying model and its results see **Table S** and Figure 4. Species: *Ac = Acer pseudoplatanus, Ae = Aesculus hippocastanum, Fr = Fraxinus excelsior, Pr = Prunus avium, So = Sorbus aucuparia, Be = Betula pendula, Ca = Carpinus betulus, Fa = Fagus sylvatica, Qu = Quercus petraea, Ti = Tilia platyphyllos*.

| *species* | *period_pairwise* | *estimate* | *SE* | *df* | *t.ratio* | *p.value* |
|---|---|---|---|---|---|---|
| *Ac* | pre - drought | -28.52 | 11.79 | 417.86 | -2.42 | **0.0423** |
| | pre - post | 11.41 | 12.03 | 422.45 | 0.95 | 0.6098 |
| | drought - post | 39.93 | 11.73 | 416.57 | 3.40 | **0.0021** |
| *Ae* | pre - drought | 20.55 | 11.67 | 415.39 | 1.76 | 0.1843 |
| | pre - post | -21.17 | 11.70 | 416.08 | -1.81 | 0.1679 |
| | drought - post | -41.72 | 11.68 | 415.54 | -3.57 | **0.0011** |
| *Be* | pre - drought | -12.19 | 12.05 | 422.74 | -1.01 | 0.5695 |
| | pre - post | 10.12 | 12.44 | 429.46 | 0.81 | 0.6949 |
| | drought - post | 22.31 | 11.74 | 416.79 | 1.90 | 0.1397 |
| *Ca* | pre - drought | 4.17 | 11.69 | 415.72 | 0.36 | 0.9323 |
| | pre - post | -28.30 | 11.74 | 416.85 | -2.41 | **0.0431** |
| | drought - post | -32.47 | 11.68 | 415.60 | -2.78 | **0.0157** |
| *Fa* | pre - drought | 32.73 | 12.21 | 418.55 | 2.68 | **0.0209** |
| | pre - post | 33.81 | 12.46 | 423.08 | 2.71 | **0.0190** |
| | drought - post | 1.08 | 12.10 | 416.39 | 0.09 | 0.9957 |
| *Fr* | pre - drought | -11.69 | 11.74 | 416.82 | -1.00 | 0.5801 |
| | pre - post | -31.73 | 11.98 | 421.56 | -2.65 | **0.0228** |
| | drought - post | -20.04 | 11.75 | 417.00 | -1.71 | 0.2043 |
| *Pr* | pre - drought | -40.64 | 11.67 | 415.35 | -3.48 | **0.0016** |
| | pre - post | 2.56 | 11.70 | 415.99 | 0.22 | 0.9740 |
| | drought - post | 43.20 | 11.68 | 415.55 | 3.70 | **0.0007** |
| *Qu* | pre - drought | 58.87 | 11.72 | 416.47 | 5.02 | **0.0000** |
| | pre - post | 63.58 | 11.78 | 417.73 | 5.40 | **0.0000** |
| | drought - post | 4.71 | 11.68 | 415.48 | 0.40 | 0.9141 |
| *So* | pre - drought | 31.84 | 11.68 | 415.63 | 2.73 | **0.0183** |
| | pre - post | 49.07 | 11.82 | 418.47 | 4.15 | **0.0001** |
| | drought - post | 17.23 | 11.73 | 416.70 | 1.47 | 0.3071 |
| *Ti* | pre - drought | 2.53 | 11.74 | 416.75 | 0.22 | 0.9748 |
| | pre - post | 6.41 | 11.80 | 418.05 | 0.54 | 0.8501 |
| | drought - post | 3.88 | 11.67 | 415.47 | 0.33 | 0.9409 |

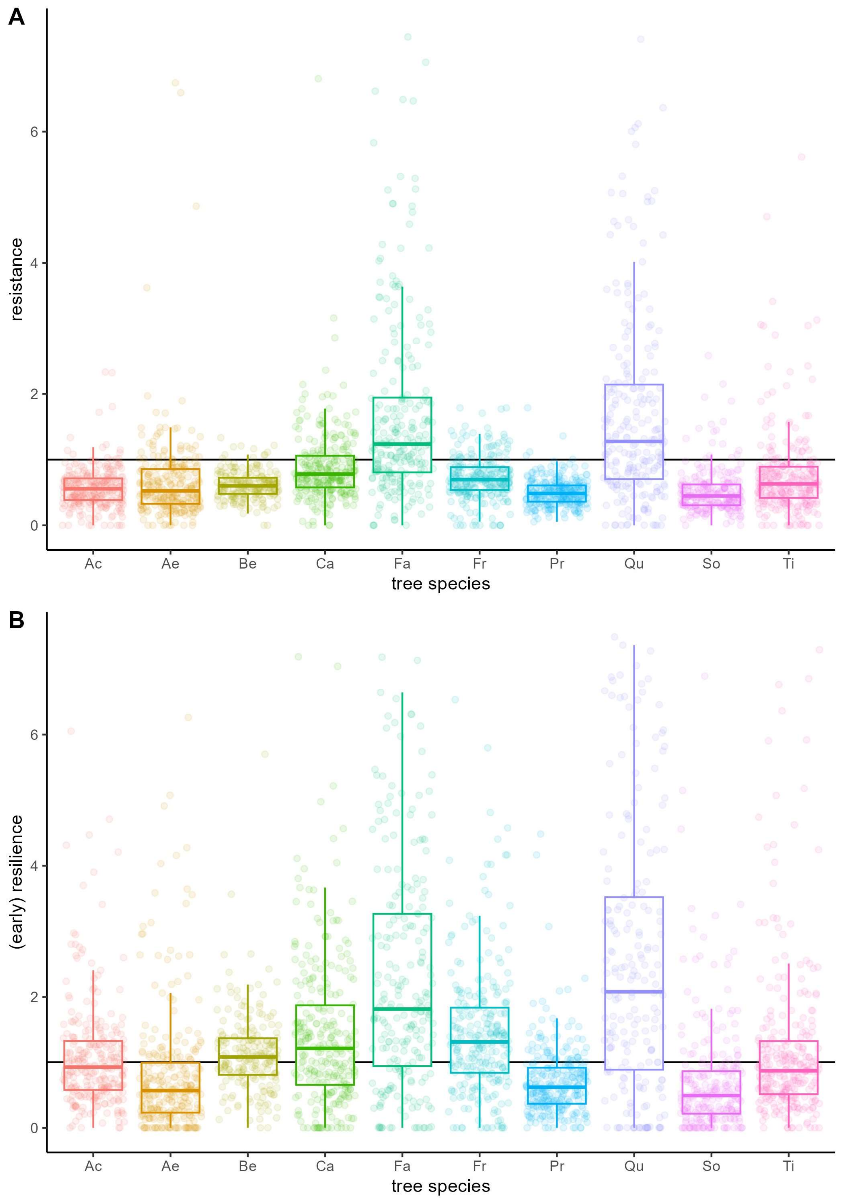

**Figure S9: Species resistance and resilience.** Resistance (A) and resilience (B) of all investigated tree individuals shown in grouped boxplots per tree species. The horizontal line at the intercept of y=1 serves as a visual support for interpretation: values on this horizontal line are tree individuals that grew as much during drought than before (resistance, A) or that grew as much after drought than before (resilience, B), values above the line stand for high resistance and resilience, i.e. tree individuals grew even more during drought than before (resistance, A) or more after drought than before (resilience, B). For better readability of the figure, 31 positive outlier for resistance (A) and 88 positive outlier for resilience (B) are not shown
